# Tumors Located in the Brain Impair the Frequency and Phenotype of Dendritic Cells in Blood and Tumor

**DOI:** 10.1101/2025.11.05.686441

**Authors:** Bryan Gardam, Tessa Gargett, Eunwoo Nam, Sidra Khan, Rebecca Jane Ormsby, Santosh I Poonnoose, Julie M Bracken, Anupama Pasam, Sakthi Lenin, Briony L Gliddon, Melinda N Tea, Chloe L Shard, Stuart M Pitson, Guillermo A Gomez, Katherine A Pillman, Shahneen Sandhu, Michael P Brown, Lisa M Ebert

**Affiliations:** Adelaide Medical School, The University of Adelaide, Adelaide, South Australia, Australia. (BG, TG, SMP, MPB, LME); Centre for Cancer Biology, University of South Australia and SA Pathology, Adelaide, South Australia, Australia. (BG, TG, EN, SK, JMB, SL, BLG, MNT, CLS, SMP,GAG, KAP, MPB, LME); Cancer Clinical Trials Unit, Royal Adelaide Hospital, Adelaide, South Australia, Australia. (TG, SK, MPB, LME); Clinical and Health Sciences, University of South Australia, Adelaide, South Australia, Australia. (TG, EN, LME); Flinders Health and Medical Research Institute, College of Medicine and Public Health, Flinders University, Adelaide, South Australia, Australia. (RJO, SIP); Department of Neurosurgery, Flinders Medical Centre, Adelaide, SA, Australia. (SIP); Department of Medical Oncology, Peter MacCallum Cancer Centre, Melbourne, Victoria, Australia. (AP); Sir Peter MacCallum Cancer Department of Oncology, University of Melbourne, Melbourne, Australia. (SS)

**Keywords:** Glioblastoma, Brain Tumor, Dendritic Cells, Immunotherapy, Anti-Tumor Response

## Abstract

**Background:** Professional antigen-presenting dendritic cells (DC) are critical for anti-tumor immune responses, yet patients with glioblastoma, an aggressive primary brain tumor that responds poorly to current investigational immunotherapies, appear to be deficient in DC. The extent of this deficiency, the specific DC subsets affected, and the causative mechanisms remain undefined. Furthermore, DCs in other brain tumors have not been systematically investigated.

**Methods:** High-parameter flow cytometry was used to profile circulating and intra-tumoral DCs in patients with glioblastoma, low-grade gliomas, and brain metastases, and non-CNS cancers. Plasma DC growth factors were quantified using ELISA. We also evaluated single-cell RNA sequencing (scRNAseq) datasets to compare intra-tumoral DCs in brain and lung tumors, and quantified DC number and phenotype in three intracranial mouse brain tumor models.

**Results:** Our studies reveal a profound systemic reduction of multiple DC subsets in the blood of patients with diverse brain tumors, coupled with reduced DC activation marker expression and lower plasma levels of FLT3L and G-CSF. Furthermore, scRNAseq analyses revealed reduced intra-tumoral DCs in glioblastoma compared to lung tumors. Circulating DC numbers inversely correlated with perioperative corticosteroid dose in patients with or without a brain tumor. However, brain tumor patients not receiving corticosteroids also had reduced DCs, suggesting a direct effect of the brain tumor. This was supported by our observation of systemic DC defects in mouse brain tumor models.

**Conclusions:** We reveal profound DC defects in patients with brain tumors, which may contribute to current difficulties in developing effective immunotherapies for glioblastoma.

**Summary:** We demonstrate multiple DC defects in patients with brain tumors. This includes a profound reduction in circulating DC number, diminished activation marker expression and growth factor levels in cancer patients with brain tumors compared to those without, and reduced intra-tumoral DCs in brain compared to lung tumors. This is the first time DC subsets have been fully characterized in a range of brain tumor patients. We show that corticosteroid usage is closely associated with DC defects, highlighting the adverse effects of a standard symptomatic treatment on these critical immune cells. However, tumors located within the brain also directly contribute to DC defects. We identified several mouse brain tumor models that can be used to further the understanding of this endogenous DC deficiency and to develop approaches to restore DCs, ultimately leading to new combination immunotherapies for the treatment of brain cancers.

**Highlights:**

- DCs are reduced in brain tumor patients and mice with intracranial tumors.
- DCs are rare within glioblastoma tumor tissue.
- The presence of a brain tumor and corticosteroid use are both associated with DC defects.

## Introduction

Glioblastoma is a devastating, aggressive form of malignant brain tumor with a 5-year overall survival rate of 6.8%^1^. The standard of care treatment has changed little over the last 20 years, highlighting the urgent need for new treatments^2^. Current immunotherapies, including chimeric antigen receptor T cells (CAR-T cells), DC vaccines, and immune checkpoint inhibitors, have all been investigated for the treatment of glioblastoma and are well summarised elsewhere^3,4^. However, significant treatment advances have not yet been made, potentially due to profound local and systemic immune suppression in these patients^4^.

Professional antigen-presenting DCs are understood to be critical in anti-tumor responses, particularly the antigen-cross-presenting classical dendritic cells type 1 (cDC1s)^5^. More recently, it has been identified that each DC subset has an important role to play in anti-tumor immune responses^6^. Classical DC type 2 (cDC2s) have often been considered to be the pro-tumor DC subset; however, recent studies have identified the important anti-tumor role they play in both the support of cDC1s and in the absence of cDC1s^7,8^. Furthermore, two subsets of cDC2s have been identified based on the expression of CD5. CD5^+^cDC2s have a greater ability to migrate to lymph nodes and produce strong regulatory T cell responses and immune regulatory interleukins. Conversely, CD5^-^cDC2s migrate more slowly but produce strong interferon-gamma (IFN-γ) T cell responses^9,10^.

The more recently identified dendritic cell type 3 (DC3s)^11,12^ are the predominant DC subset in melanoma, breast and lung cancers^13–16^. While some have considered that DC3s and monocyte-derived DCs are the same, DC3s have been described as a discrete DC subset by others^11,17,18^. A recent study has shown that DC3s can secrete high levels of pro-inflammatory cytokines, but they have limited ability to stimulate cytotoxic T cell responses^19^. In contrast, an earlier study showed that DC3s provide essential support to cytotoxic T cells by trans-presenting IL-15^20^. Recent reports also demonstrate the need for effective DCs in immune checkpoint inhibitor (ICI) therapies^21^. DCs have therefore been identified as being critical for providing cytotoxic T cells with proliferation and survival signals in the tumor microenvironment (TME)^20–22^.

Important growth factors for DC development include FMS-like tyrosine kinase 3 ligand (FLT3L), which is the critical growth factor required for cDC1s and cDC2s, whereas granulocyte-macrophage colony-stimulating factor (GM-CSF) has been identified as the required growth factor for DC3 development, independent of FLT3L^18,23^. In contrast, granulocyte colony-stimulating factor (G-CSF), has been identified as an inhibitor of cDC1 differentiation^18,24^, although it is an important growth factor that can mobilize cells from the bone marrow^25^.

In appreciating the vital role DCs play in immune responses to cancer, DCs have been reported to be reduced in number and less functional in the TME and peripheral blood of glioblastoma patients^26–31^. These previous studies have considered different markers and DC subsets, ranging from total DCs and plasmacytoid DCs (pDCs)^27^ to cDC2s and pDCs^28,29^. Only two studies looked at cDC1s, with one reporting a reduction of 40%^30^ and the other reporting no DCs identified^31^ in peripheral blood. All of these studies only considered circulating DCs. Further, these studies used a variety of preparations for their observations, including fresh whole blood^27,28,30^, DCs purified from peripheral blood^31^, or expressed as the percentage of CD45^+^ cells^26^, making them difficult to compare. However, no study thus far has directly compared all DC subsets, considered brain tumors beyond glioblastoma, or compared DCs within brain tumors to other more immunogenic tumors. As such, a complete picture of the DC subsets in patients with brain tumors is lacking. As DCs are required for effective T-cell mediated control of tumors, DC deficiency represents a significant hurdle to immune-based therapy and requires further understanding, including its mechanistic basis.

In this study, we explored all currently defined DC subsets in primary and recurrent glioblastoma patients, patients with low-grade gliomas (LGG), patients with brain metastasis, and patients with other cancers not affecting the brain to investigate changes in DC numbers and functional markers. We then extended our findings via analysis of scRNAseq datasets and intracranial mouse models.

## Results

### Brain tumor patients exhibit significant reductions in circulating DCs compared to healthy donors or cancer patients without a brain tumor

Using high-parameter flow cytometry adapted from Mair and Liechti^32^ to identify DC subsets, we analyzed PBMCs from healthy donors (n=17) and patients with histologically confirmed primary glioblastoma (n=14), recurrent glioblastoma (n=16), LGG (n=7), melanoma and NSCLC metastases to the brain (n=22), and patients with metastatic melanoma and NSCLC not involving brain (non-brain tumor) (n=17). Median age varied between some groups (Fig 1A). Full details of the participants in this study are in Table S1.

**Figure 1.**
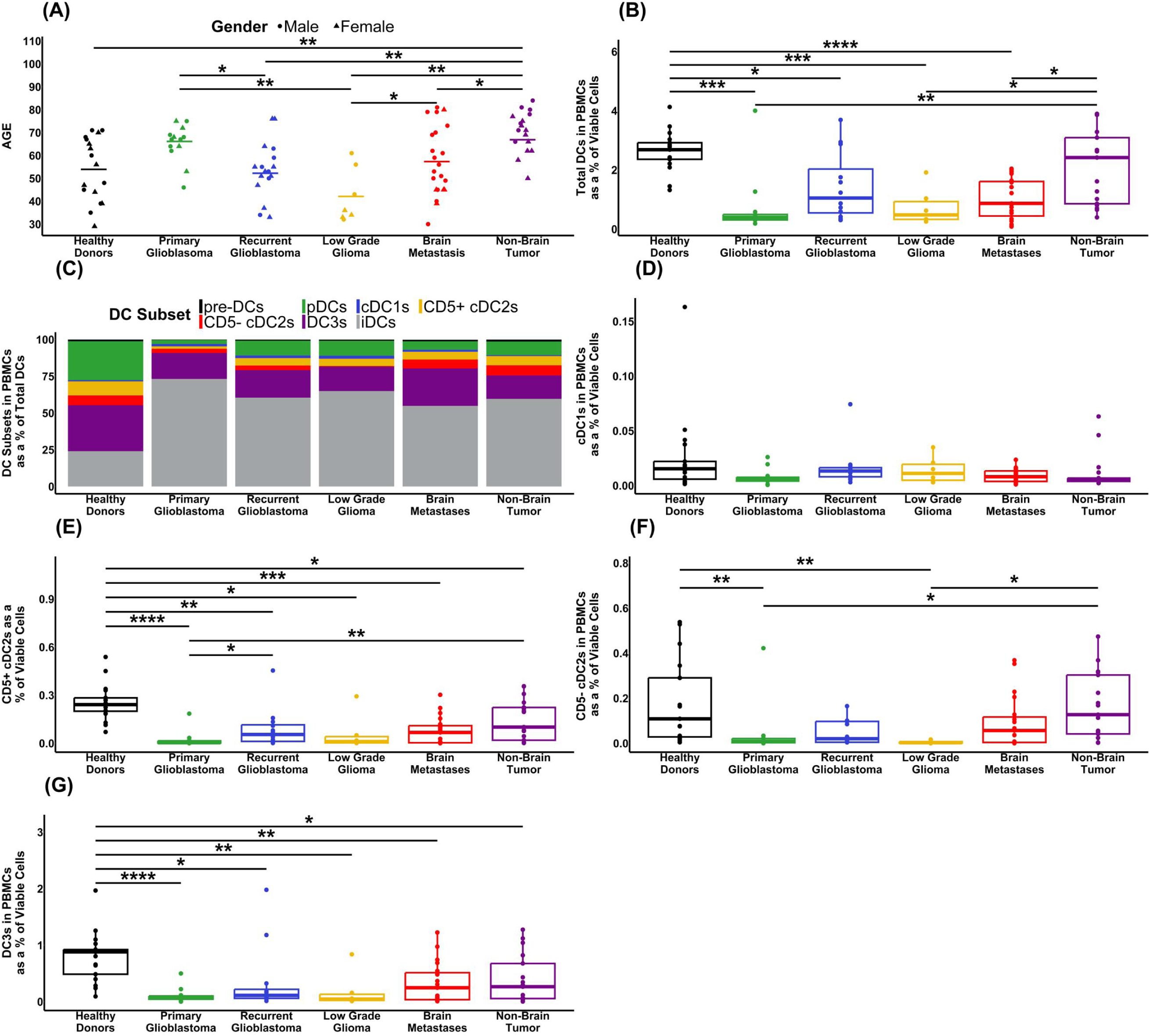
Patient demographics and analysis of their circulating DCs by flow cytometry. (A) Summary of the healthy donors and patients in this study by age, gender and diagnosis. (B) Frequency of total DCs (viable HLA-DR^+^ CD3^-^ FcεR1a^-^ CD19^-^ CD20^-^ CD88^-^ CD89^-^ cells) as a percentage of total viable mononuclear cells. (C) DC subsets as a percentage of total DCs, defined as pre-DCs (CD123^+^ CD5^+^), pDCs (CD123^+^ CD5^+^), iDCs (CD123^-^ XCR1^-^ CD11c^+/-^ FcεR1a^+/-^ HLA-DR^Low^ CD86^-/Low^). (D) cDC1s (CD123^-^ XCR1^+^); (E) CD5^+^ cDC2s (CD123^-^ XCR1^-^ CD11c^+^ FcεR1a^+^ CD5^+^); (F) CD5^-^ cDC2s (CD123^-^ XCR1^-^ CD11c^+^ FcεR1a^+^ CD5^-^); (G) DC3s (CD123^-^ XCR1^-^ CD11c^+^ FcεR1a^+^ CD5^-^ CD163^+^ CD14^+/-^). (D-G) Percentages of DC subsets across the patient groups, as a percentage of total mononuclear cells.

We identified a significant reduction in the total circulating DCs (indicative gating Fig S1A) for all patients with a brain tumor compared with healthy donors. (Fig 1B). The most dramatic difference was observed in primary glioblastoma patients, who had a 6.75-fold reduction in median DC frequency relative to healthy donors. When compared to melanoma or NSCLC patients without a brain tumor, most other brain tumor groups also had a significant reduction in the proportion of DCs, with the exception of recurrent glioblastoma patients where the level of inter-patient variability was greatest. There was no significant difference in the total circulating DCs between the healthy donors and non-brain tumor patients (Fig 1B).

We then investigated the individual subsets of circulating DCs (indicative gating is at Fig S1B) and identified the proportional changes in each subset (Fig 1C). While analysis of the total circulating DCs indicated no significant change between healthy donors and non-brain tumor patients, this more detailed analysis revealed a shift in subset frequencies. Of interest, immature DCs (iDC)^33^ dominated the total DC population in all cancer patients compared to healthy donors, and plasmacytoid DCs (pDCs) and DC3s were markedly reduced in all cancer patients, irrespective of tumor type.

In further analysis of each DC subset as a percentage of total viable PBMC we observed no significant change in cDC1s between any groups (Fig 1D). In contrast, the CD5^+^ subset of cDC2s was significantly reduced in all groups of patients compared to healthy donors (Fig 1E). The CD5^-^ subset of cDC2s showed a significant reduction only in primary glioblastoma and LGG compared to healthy donors, with no difference observed between non-brain tumor patients and healthy donors (Fig 1F). DC3s showed a significant reduction in patients of all groups compared to healthy donors (Fig 1G).

We also assessed additional DC subsets that do not play a clearly identified role in anti-tumor immunity (Fig S2). Precursor DCs (pre-DCs) displayed a significant reduction across all brain tumor groups (but not cancer patients without brain tumors) compared to healthy donors (Fig S2A). On the other hand, we identified a significant increase in non-brain tumor patients’ iDCs compared to primary glioblastoma, LGG and brain metastasis patients (Fig S2B). All groups had a significant reduction in circulating pDC compared to the healthy donors (Fig S2C).

In summary, immature DCs dominate in cancer patients compared to healthy donors, while patients with brain tumors had decreases in both total DCs and in all mature DC subsets, except for the rare cDC1s, compared to healthy donors. Strikingly, and in contrast to brain tumor patients, patients with non-brain tumors had no significant change in total DC frequency compared to healthy donors, with differences only observed in the CD5^+^ cDC2s and DC3 subsets.

### Circulating levels of key DC growth factors are perturbed in brain tumor patients

Next, we measured by ELISA the plasma levels of growth factors that may contribute to a reduction in DCs. There was a significant reduction in FLT3L levels in patients with primary glioblastoma, LGG and brain metastases, compared to both healthy donors and non-brain tumor patients (Fig 2A). We also assessed the levels of G-CSF, and while fewer of the samples had detectable levels, there was a significant reduction in primary glioblastoma, LGG and brain metastasis patients compared to healthy donors or non-brain tumor patients (Fig 2B). We then assessed the levels of GM-CSF in the plasma, and observed a trend toward increased plasma GM-CSF in patients with brain tumors compared to healthy donors. Interestingly, none of the non-brain tumor patients had detectable levels of GM-CSF, in contrast to patients with primary glioblastoma and LGG (Fig 2C). Thus, there are clear differences among brain tumour patients in circulating levels of DC growth factors compared to healthy donors and non-brain tumor patients, suggesting that these factors may be involved in the pathogenesis of DC deficiencies in brain tumor patients.

**Figure 2.**
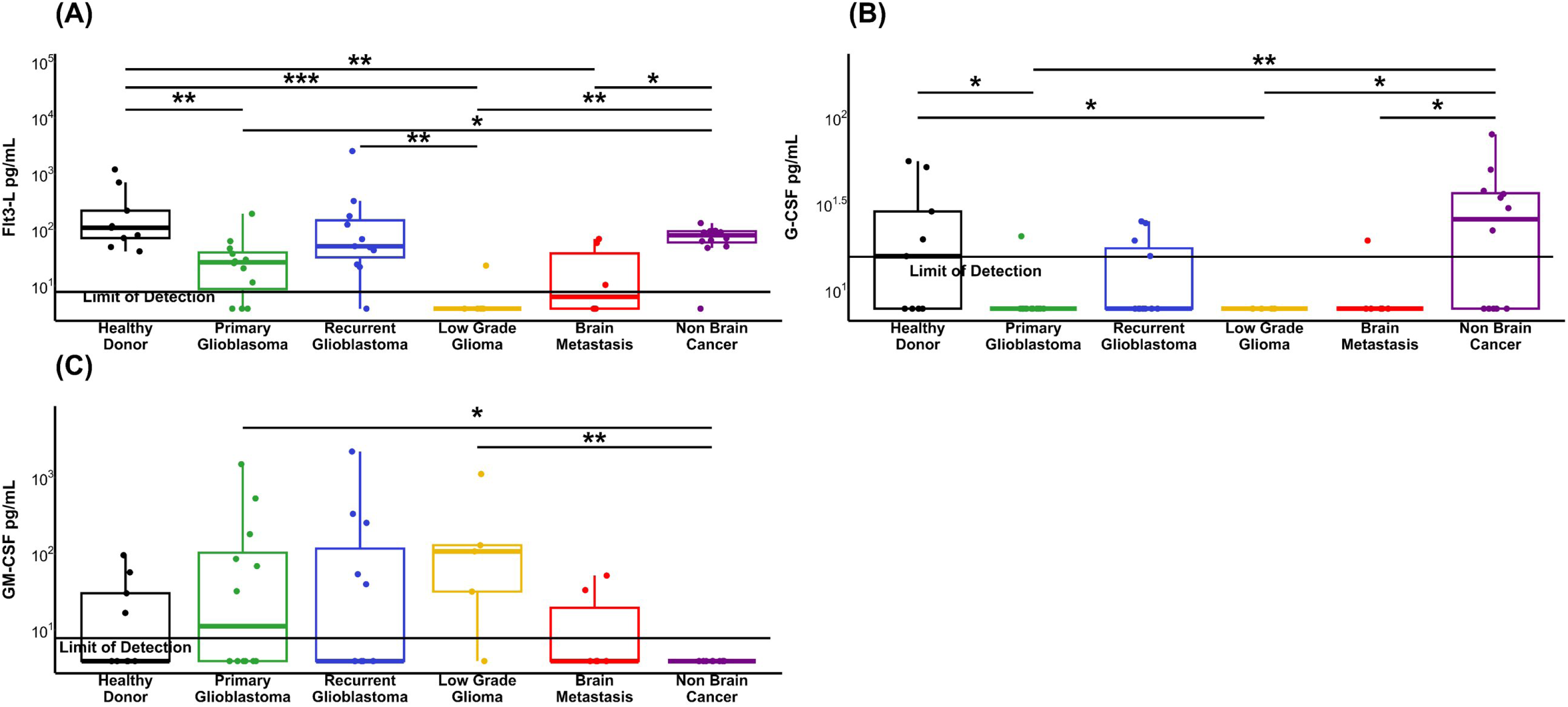
Quantification of DC growth factors in patient plasma (A) Quantification of FLT3L in patient plasma. (B) Quantification of G-CSF in patient plasma, and (C) Quantification of GM-CSF in patient plasma. Results below the limit of detection (LOD) were calculated as half of the lower LOD as described in Giskeødegård & Lydersen^53^.

### Patients with brain tumors have reduced DC activation marker expression

We next investigated the expression levels of key functional markers on DC subsets by comparing the mean fluorescence intensity (MFI) between patient groups and healthy donors. We focused these analyses on HLA-DR, CD86 (markers of DC activation and T cell engagement), and PD-L1 (immune inhibitory marker). Patients with primary glioblastoma showed a significant decrease in HLA-DR expression in their cDC1 subset compared to healthy donors (Fig 3A). Significant decreases in HLA-DR expression were also observed in CD5^+^ (but not CD5^-^) cDC2s from primary glioblastoma patients compared to healthy donors (Fig 3B-C). The DC3s showed a significant reduction in HLA-DR expression for primary glioblastoma, brain metastases and non-brain tumor patients compared to healthy donors (Fig 3D). The T cell costimulatory marker CD86 demonstrated a significant increase of expression in cDC1s from non-brain tumor, brain metastases, and primary glioblastoma patients compared to healthy donors (Fig 3E). In contrast, CD86 expression in CD5^+^ DC2s was significantly reduced for all brain tumor patients except for LGG compared to healthy donors (Fig 3F), with a similar pattern of reduction in CD5^-^ cDC2s and DC3s (Fig 3G-H). Finally, we examined the T cell inhibitory ligand PD-L1 and found it was elevated on multiple DC subsets from non-brain tumor patients compared to most other groups. (Fig 3I-L, Fig S3C). Of interest, the non-brain tumor patient cohort consists of melanoma and NSCLC patient groups known to have high response rates to PD-1 and PD-L1 inhibitors^34^. We also observed a significant increase in PD-L1 expression in primary glioblastoma and brain metastasis patients compared to healthy donors within the cDC1 subset (Fig 3I). The expression of each marker on pre-DCs, iDCs, and pDCs is shown in Fig S3.

**Figure 3.**
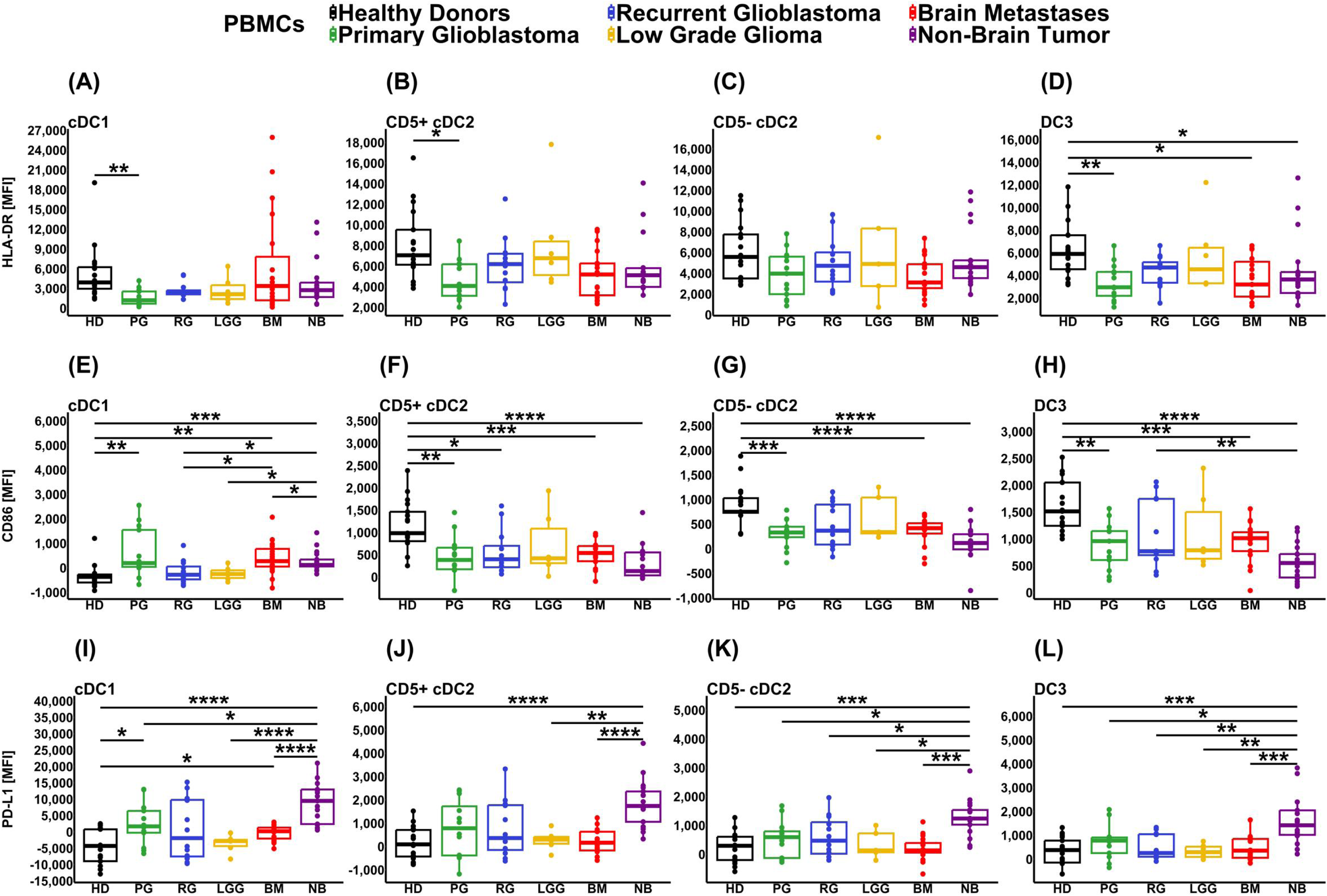
Quantification of circulating DC functional marker expression by flow cytometry. DC subsets were identified as in Figure 1, with levels of the indicated functional markers expressed as mean fluorescence intensity (MFI). Graphs show the expression of (A-D) HLA-DR, (E-H) CD86, and (I-L) PD-L1 within cDC1s (first column), CD5^+^ cDC2s (second column), CD5^-^ cDC2s (third column), and DC3s (last column). Patient groups are abbreviated as follows: Healthy donors (HD), Primary Glioblastoma (PG), Recurrent Glioblastoma (RG), Low Grade Glioma (LGG), Brain Metastases (BM), and Non-Brain Tumor (NB).

In summary, we observed lower activation marker expression in multiple DC subsets from patients with brain tumors compared to healthy donors, whereas the main difference in DC phenotype in cancer patients without a brain tumor was higher levels of PD-L1.

### Corticosteroids have a dose-dependent effect on DC frequency in patients with and without brain tumors

Corticosteroids are used routinely for symptom management in patients with brain tumors. While deleterious effects of corticosteroids on the immune system, including lymphocyte trafficking and immune-mediated killing, are well-characterized elsewhere^35^, there are also reports of brain tumors themselves directly influencing systemic immune compartments, including the loss of major histocompatibility complex class II (MHCII) expression^36^ and T cell sequestration in the bone marrow^25^. Therefore, we sought to delineate the influence of brain tumors and corticosteroids on DCs.

In line with Figure 1, we found a significant reduction in total circulating DC frequency (at the time of surgery) in brain tumor patients receiving corticosteroids either long term or in the days prior to surgery when compared to either healthy donors or non-brain tumor patients not receiving corticosteroids (Fig 4A). A less dramatic, but still significant, reduction in DCs was also observed for brain tumor patients who received corticosteroids perioperatively on the day of surgery only. Importantly, we also observed a significant reduction in DCs in brain tumor patients who did not receive any corticosteroids prior to blood collection, when compared to either healthy donors or non-brain tumor patients not receiving corticosteroids, implying a direct effect of brain-localized tumor growth on DC frequency. In contrast, total DCs were significantly reduced in non-brain tumor patients treated with corticosteroids to manage concurrent conditions, when compared to those that received no corticosteroids or healthy donors. This highlights a likely role for corticosteroids in reducing DCs regardless of tumor location. Indeed, within the entire cancer patient cohort, there was a strong correlation between DC frequency and the total dose of corticosteroids given prior to blood collection (Fig 4B).

**Figure 4.**
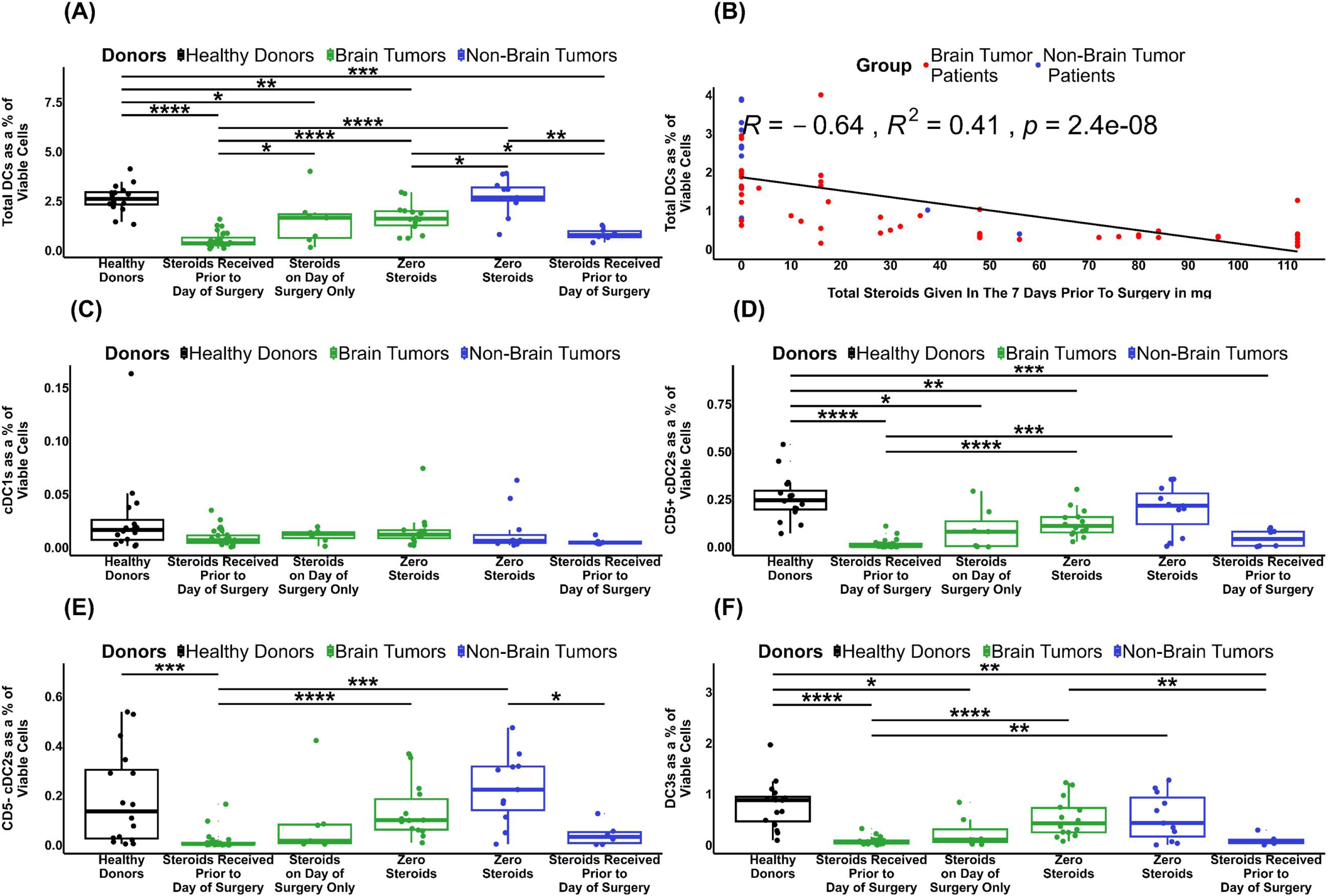
Association between circulating DC frequency and steroid dosage. (A) total DCs in patients that received corticosteroids in the days leading up to surgery, on the day of surgery only, and patients not receiving corticosteroids, compared to healthy controls. (B) correlation of total DCs versus total corticosteroids given in the 7 days prior to surgery. (C-F) differences in circulating DC subsets between patients receiving or not receiving steroids: (C) cDC1s, (D) CD5^+^ cDC2s, (E) CD5^-^ cDC2s, and (F) DC3s.

When considered as DC subsets, we observed no significant difference in cDC1s between any groups (Fig 4C). However, both CD5^+^ and CD5^-^ cDC2s were reduced in brain tumor patients receiving corticosteroids prior to surgery compared to healthy donors or non-brain tumor patients not receiving corticosteroids. (Fig 4D-E). Finally, the DC3s showed a significant reduction in brain tumor patients receiving corticosteroids prior to surgery, and in brain tumor patients who received corticosteroids on the day of surgery, compared to healthy donors (Fig 4F). Pre-DCs, iDCs, and pDCs were also significantly reduced in brain tumor patients receiving corticosteroids compared to healthy donors, and compared to non-brain tumor patients not receiving corticosteroids (Fig S4A-C). Interestingly, we observed a significant increase in iDCs among non-brain tumor patients not receiving corticosteroids compared to healthy donors (Fig S4B).

In summary, steroid dose was closely linked to reduced DCs in both brain tumor and non-brain tumor patients. In general, all the DC subsets, except for cDC1s, mirrored the observations of the total DCs. At the same time, patients with brain tumors not receiving steroids had fewer DCs than healthy donors or patients with non-brain tumors, suggesting a tumor location-linked mechanism.

### Brain tumors have fewer DCs in the TME compared to lung tumors

Next, we extended our investigation to include analysis of brain tumor tissues. To achieve this, we used publicly available scRNA-seq datasets to determine the frequency of DCs within the TME of patients with glioblastoma, and compared these to NSCLC tumors, which are generally considered immune ‘hot’ (Fig 5). DC-enriched cell clusters (Fig 5A-B and Fig S5) were iteratively subsetted from mixed population cell clusters (Fig S5) using a curated list of broad cell type and DC subset-specific marker genes (Fig 5C-D, Fig S5, Table S2). High expression of C5AR1, a monocyte marker not typically expressed by cDC2 or DC3 cells was used to distinguish DCs from monocytes within the myeloid lineage cluster^37^. Using this approach, we observed a striking and statistically significant decrease in the total DCs as a percentage of immune cells in both primary and recurrent glioblastoma compared to lung cancer (Fig 5E). Cells were classified as immune cells if more than 70% of clustered cells expressed PTPRC (CD45). A significant decrease was also observed in cDC1s, cDC2s, and pDCs in primary glioblastoma compared to non-brain tumor patients (Fig 5F-G, I), with only DC3s showing no significant changes (Fig 5H).

**Figure 5.**
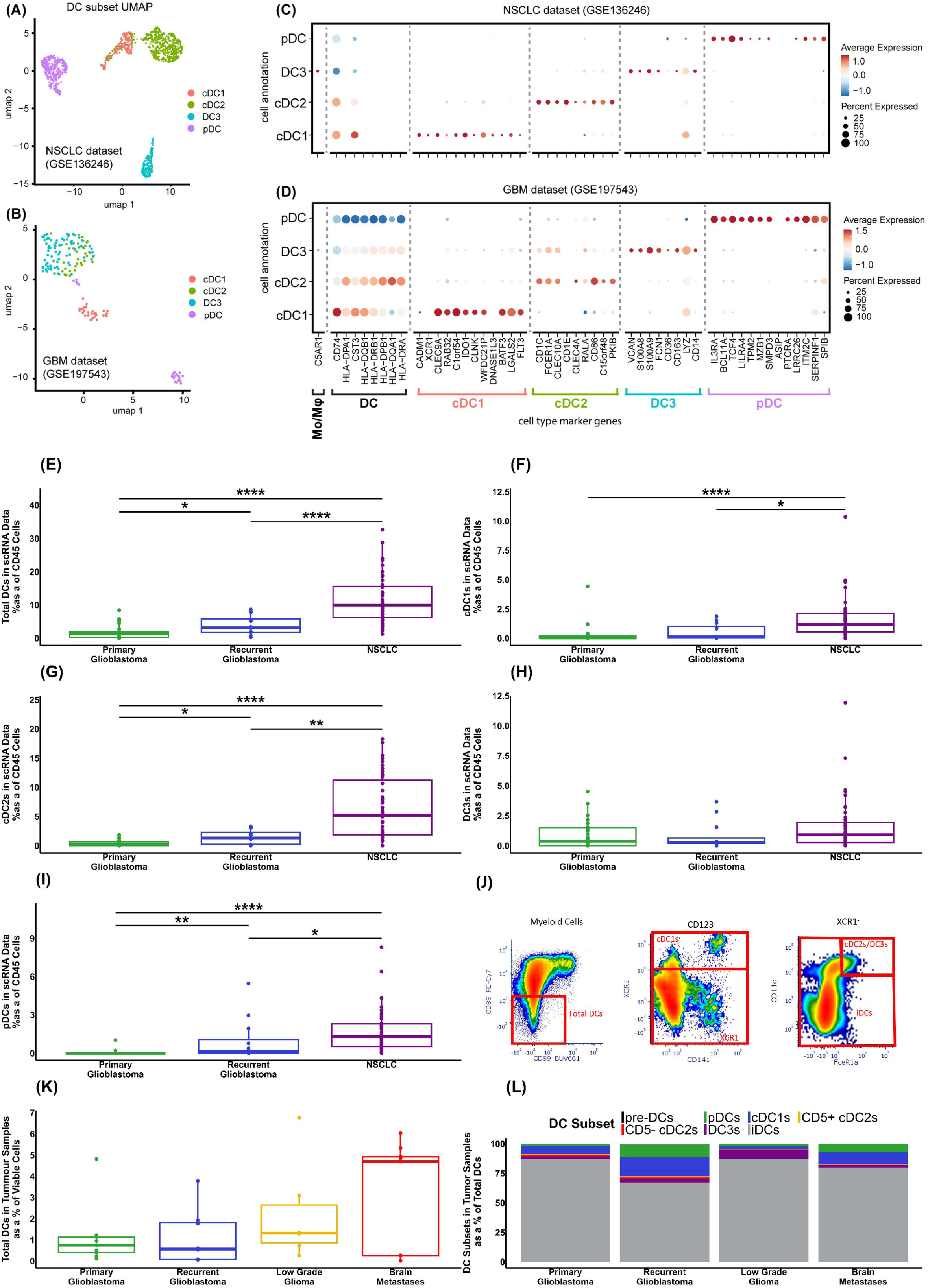
Quantification of DCs in tumor tissue using publicly available scRNA-seq data for primary glioblastoma, recurrent glioblastoma, and lung cancer patients. (A, B) Representative UMAP plots showing DC subset annotation in the NSCLC (GSE136246) and glioblastoma (GSE197543) datasets, respectively. (C, D) Corresponding marker gene expression for DC subsets in the NSCLC and glioblastoma datasets. UMAP plot of the other datasets is shown in Figure S5C. (E-I) Cell numbers identified in scRNAseq data sets are shown as a percentage of CD45^+^ cells: (E) Total DCs, (F) cDC1s, (G) cDC2s, (H) DC3s, and (I) pDCs. (J) Indicative flow cytometry gating of DCs in patient tumor samples, Myeloid cells identified as CD3^-^, HLA-DR^+^, CD19^-^, CD20^-^, (K) Quantification of total DCs identified by flow cytometry in patient tumor samples, and (L) DC subsets identified in patient tumor samples.

We also confirmed the presence of these diverse DC subsets in dissociated patient brain tumor samples using flow cytometry. In line with the scRNAseq analyses, we were able to identify all of the DC subsets by flow cytometry (Fig 5J), using the same gating strategy as for PBMCs (Fig S1). DCs constituted a small proportion of total viable cells (range 0.02 – 6.73%), with no significant differences among patients from different brain tumor groups (Fig 5K). In all groups, the majority of the identified DCs were iDCs, although other subsets were detectable in varying proportions (Fig 5L). Finally, we considered the functional markers expressed by DCs within brain tumors compared to healthy donor PBMCs. The most striking changes were observed for CD86, which was increased on cDC1 from all brain tumors compared to PBMC, yet paradoxically decreased on DC3s within tumors (Fig S6). In addition, CCR7 was reduced on cDC1 within tumors, as may be expected for tissue-resident cells.

These multimodal analyses confirm, for the first time, the presence of diverse DC subsets within the brain tumor microenvironment. However, the frequency of DCs in brain tumors was dramatically reduced compared to lung tumors, highlighting the stark differences between the typically cold tumor immune microenvironment of brain cancer patients and the hot tumor immune microenvironment of lung cancer patients.

### Murine models of brain tumors replicate the DC defects seen in patients

Our analysis indicated that corticosteroid use is closely associated with the loss of DCs observed in brain cancer patients. However, brain tumor patients not receiving corticosteroids also demonstrated a significant reduction in DC frequency. To confirm that brain tumors directly contribute to systemic DC deficiency, we examined splenic DCs in three orthotopic brain tumor mouse models generated by the intracranial engraftment of CT2A, GL261 or SB28 murine glioma cell lines into C57BL/6 mice, in the absence of systemic corticosteroids. All glioma cell lines were engineered to express the disialoganglioside GD2 by stable expression of murine GD2 and GD3 synthases^38^. GD2, which is identical between mice and humans and expressed at low level in normal brain of both species, is a target antigen of interest in brain cancer CAR-T cell development^39^. We used flow cytometry to analyze splenic DCs, with the indicative flow cytometry gating shown in Fig S7. We found a significant reduction in total DCs in the spleens of mice bearing SB28 and CT2A tumors compared to control C57BL/6 mice (Fig 6A), and while not significant, a trend towards a reduction in total DCs in the spleens of mice bearing GL261 tumors compared to control mice. We then explored DC subsets and identified a reduction in cDC1s and iDCs, with a concurrent increase in cDC2s and DC3s, in the spleens of all tumor models compared to control mice (Fig 6B-E). Finally, we examined the expression of functional markers. There was a significant reduction in MHC II expression in all DC subsets of mice bearing CT2A tumors compared to healthy mice, although this trend was reversed for mice bearing SB28 tumors (Fig 6F). For CD86 expression, we observed a significant reduction in all DC subsets of the CT2A tumor-bearing mice compared to healthy mice (Fig 6G). Finally, for PD-L1 expression, we saw a significant reduction in expression on cDC2s of CT2A tumor-bearing compared to healthy mice (Fig 6H). Therefore, even in the absence of steroids, DC numbers were significantly reduced in the spleens of two out of three intracranial models of glioblastoma, and the expression of CD86 was also reduced, suggesting a reduced ability of DCs to co-stimulate T cells.

**Figure 6.**
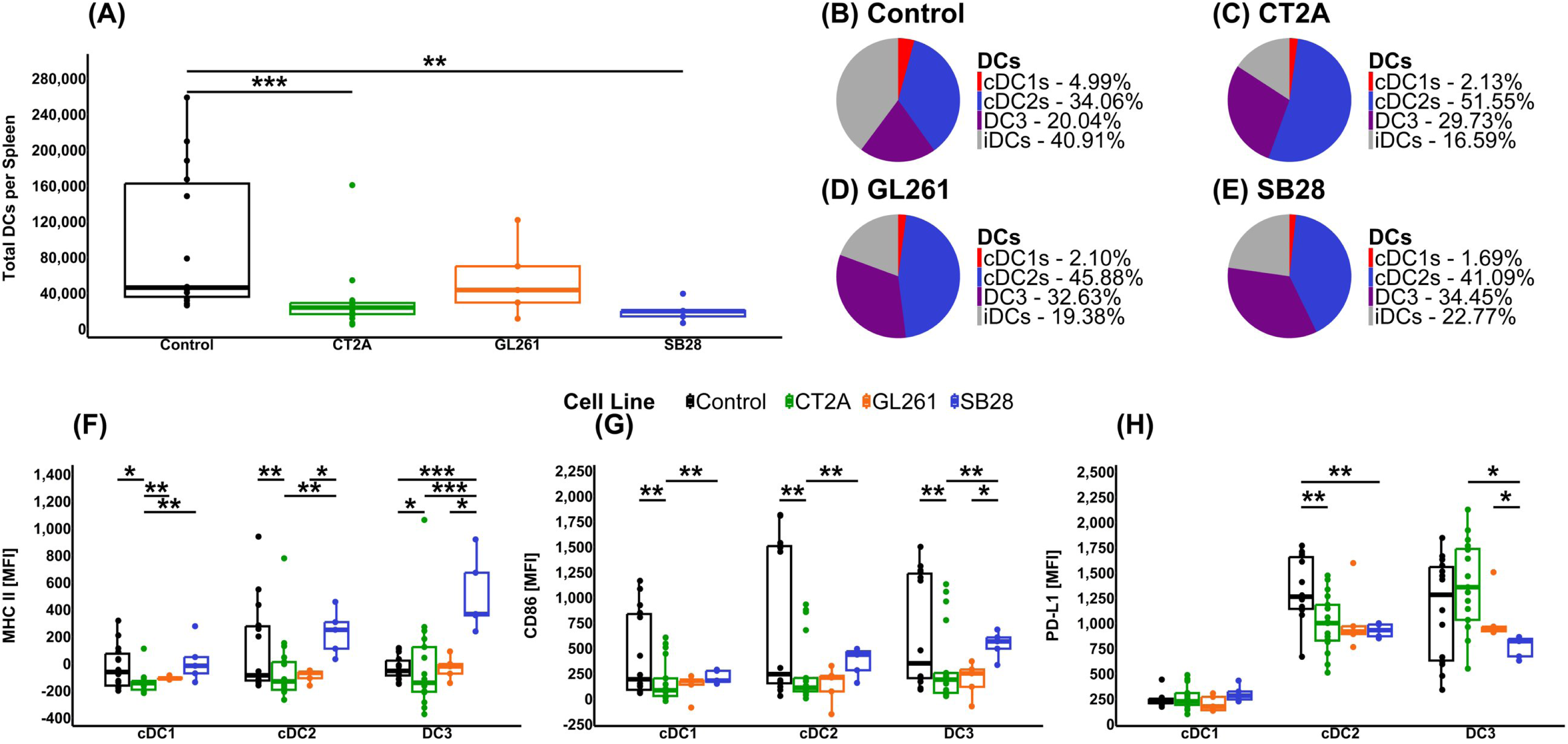
Comparison of spleen DC populations in orthotopic immunocompetent murine models of GD2 expressing brain tumors. (A) Absolute number of DCs in the spleens of C57Bl/6 mice with or without the indicated brain tumors, as determined by flow cytometry using TruCount tubes for absolute cell enumeration. (B-E) Pie charts demonstrate DC subsets as a percentage of total DCs from the spleens of (B) control mice, (C) CT2A tumor-bearing mice, (D) GL261 tumor-bearing mice, (E) SB28 tumor-bearing mice, (F-I) Functional markers for cDC1, cDC2, and DC3 from mouse spleens; (F) MHC II, (G) CD86, (H) PD-L1.

## Discussion

Although select DC populations have been shown by others to be reduced and less functional in patients with glioblastoma^26–30^, here we have shown a reduction in all circulating DC subsets, except cDC1s, across patients with glioblastoma, low-grade gliomas, and brain metastases. In contrast, cancer patients with tumors outside the brain had no overall reduction in total DCs and displayed less extensive changes in DC subsets. The greatest inter-patient variation was observed among recurrent glioblastoma patients who varied in clinical factors such as time to recurrence and treatment stage. The extent of this variation does limit interpretation of data in this clinical cohort.

When we examined plasma growth factor levels, we saw a significant reduction in FLT3L in brain tumor patients compared to both healthy donors and non-brain tumor patients. As the major growth factor for the production of classical DCs^18,23^, this could account for the reduction in cDC2s. High levels of G-CSF can be associated with reduced classical DCs^18,24^. Our results showed that brain tumor patients generally had lower levels of G-CSF compared to healthy donors and non-brain tumor patients, which would not explain the reduction in classical DCs. In contrast, GM-CSF has been identified as a key growth factor for generating DC3s^11,18^. Levels of GM-CSF were generally higher in the plasma of glioblastoma and low-grade glioma patients than in the healthy donors and non-brain tumor patients; however, we did not observe a corresponding increase in the number of DC3s in these patients. Meanwhile, GM-CSF has been shown to have an immunosuppressive effect in glioblastoma^40^ and is increased following traumatic brain injuries^41^. In contrast, differences in levels of G-CSF and GM-CSF in brain tumor patients were less consistent and are insufficient to explain the observed changes in DCs. In particular, we had anticipated, but did not observe, a decrease in GM-CSF, which has been reported as a key growth factor for generating DC3s^11,18^. Our findings, therefore, suggest that DC3 generation involves additional essential growth factors.

When considering markers of DC function, we observed a decrease in HLA-DR expression in cDC1s, CD5^+^cDC2s, and DC3s, and a decrease in CD86, in most DC subsets in primary glioblastoma. This suggests that, in addition to a reduction in frequency, the antigen presentation and T cell co-stimulatory capacity of residual DCs in brain tumor patients is reduced compared to healthy donors. We also observed an increase in the inhibitory ligand PD-L1 on cDC1s in brain tumor patients compared to healthy donors, suggesting that the ability of cDC1 to activate T cells is compromised in these patients, even though the frequency of this subset was unaffected. Taken together, this suggests that DCs in primary glioblastoma patients are of a less activated phenotype, with impaired ability to stimulate anti-tumor T cells, compared to the same subsets from healthy donors. However, we have not conducted functional studies on the DCs to determine if these phenotypic differences translate into a reduced ability to stimulate T cells, and this remains an area for further research.

Although corticosteroids are important for symptomatic treatment of brain tumor patients, the adverse effects of steroids on the immune system are well-reported^42–46^. A recent study by Dusoswa et al. reported systemic immune suppression in glioblastoma patients, and the immunosuppressive effects of steroids, including reductions in CD4^+^ T cells, alternative and intermediate monocytes, with increased B cells, NK cells and double negative T cells. Interestingly, these authors also reported an increase in B cells and DCs in glioblastoma patients, but this difference was lost when corrected for age and sex, suggesting that age played a significant role^47^. A reduction in CD4^+^ T cells in patients receiving corticosteroids has also been reported by others^48^. In addition to showing that the cumulative dose of administered steroids inversely correlates with the number of DCs, we also reveal that the presence of a brain tumor alone (in the absence of steroid treatment) is associated with reduced numbers of circulating DCs. These data are consistent with previous reports of reduced circulating T cell numbers in brain tumor patients and brain tumor-bearing mice, which are attributed to brain-tumor provoked peripheral immune suppression^25^. This phenomenon, independent of steroid usage and due to systemic release of unknown factors resulting from tumor-mediated brain injury, has been recognized by others^25,49,50^. Our own investigations in three orthotopic mouse models support a direct, brain-tumor mediated suppression of DC number and function. In our models, we observed a significant reduction in DC numbers in the spleens of two of our models, coupled with significant reductions in the expression of MHC II and CD86 in the CT2A model, similar to our observations in primary glioblastoma patients.

We also used scRNAseq data to examine the number of DCs in the TME of primary and recurrent glioblastoma tumors, and compared these with scRNAseq data from NSCLC patients without brain metastases. Although the reduced numbers of circulating DCs may be important, DCs in the TME are critical for anti-tumor immune responses, through the priming, activation, and provision of proliferation and survival signals to T cells^20–22^. In all of the DC subsets examined, there were significantly fewer DCs in the brain tumor patients compared to the lung cancer patients, with the lowest in primary glioblastoma patients. The only exception to this was the DC3 subset, which nevertheless displayed a corresponding non-significant trend. Flow cytometry studies confirmed that diverse DC subsets were detectable, but very rare, within dissociated patient brain tumor specimens, and had an overwhelmingly immature phenotype. Likewise, Pombo Antunes et al. detected multiple DC subsets in the tumors of glioblastoma patients and the brains of mice bearing GL261 tumors, although there was no comparison to other cancers^26^. Thus, for the first time, we show that reduced numbers of circulating DCs in brain cancer patients compared to non-brain tumor patients also extends to the tumor microenvironment.

In this study, we have shown that patients with brain cancer have fewer DCs with a less functional phenotype in the blood and tumor compared to patients with tumors located outside the brain. While tumor-intrinsic effects likely contribute to the observed DC defects, corticosteroid use may greatly exacerbate them. As DCs are critical for T-cell mediated anti-tumor immune responses^6–8,19^, our findings strongly support future research focused on strategies to restore DC number and function in brain cancer patients, and to consider, where possible, alternatives to corticosteroids for symptom management. This could include consideration of steroid alternatives, such as the anti-VEGF agent bevacizumab^51^. Together, these approaches are expected to accelerate the successful development of immunotherapy for brain cancer patients.

## Resource Availability

### Lead contact

Requests for further information should be directed to the lead contact, Associate Professor Lisa M. Ebert

### Data availability

Data will be made available upon reasonable request; publicly available scRNAseq data was sourced from https://figshare.com/collections/An_integrated_single-cell_transcriptomic_dataset_for1_non-small_cell_lung_cancer/6222221/3, GSE163120, GSE197543, GSE182109, and GSE173278. ScRNAseq from Ebert et al.^52^, can be requested from LME and GAG.

### Ethics

All human research was performed in accordance with the principles of the Declaration of Helsinki, and was approved by the Central Adelaide Local Health Network Human Research Ethics Committee (CALHN HREC approval # R20160727), Southern Adelaide Clinical Human Research Ethics Committee (SA HREC 286.10), and the Peter MacCallum Centre Human Research Ethics Committee (approval numbers 07/38, 11/105, and 16/153); all participants provided informed written consent. For animal studies, experiments were conducted under a protocol approved by the University of South Australia Animal Ethics Committee (#U08-23).

## Acknowledgments

This project was funded by Cancer Australia Priority-Driven Collaborative Cancer Research Scheme and The Kids’ Cancer Project (Grant ID 2020344), Mark Hughes Foundation, Ray & Shirl Norman Cancer Research Foundation and NeuroSurgical Research Foundation. The authors acknowledge the support received from the NeuroSurgical Research Foundation (NRF). BG also acknowledges the support received through the NRF-Strong Enough to Live scholarship. MNT acknowledges the support of a Fay Fuller Foundation Fellowship. LME acknowledges the support of the Ray & Shirl Norman Cancer Research Trust and James & Diana Ramsay Foundation. GAG acknowledges the support of a McCleary Murchland Fellowship, grants from the NHMRC (Ideas grant 2021/GNT2013180 (GAG and LME), and 2023/GNT2030541 (GAG and CLS), the Charlie Teo Foundation Rebel Grant, and The MAWA Trust to GAG and CLS); Tour de Cure Grants (GAG, SP, and LE). CLS also acknowledges the support of an NRF Chris Adams award. The authors acknowledge the support and generosity of the patients and medical and technical staff from SA Pathology. Glioblastoma patient tissues were received from the South Australian Neurological Tumor Bank (SANTB), which is supported by Flinders University, The University of South Australia, and the NRF, which organizations made possible the collection of blood and tissue specimens. The authors wish to thank Melanoma Research Victoria and acknowledge the MRV site contributing to this work: Peter MacCallum Cancer Centre. The authors also acknowledge the support of the Adelaide Health and BioMedical Precinct Cytometry Facility SAHMRI, the Facility is generously supported by the Detmold Group, McMahon Family, Australian Cancer Research Foundation, Cancer Council and the Australian Government through the Zero Childhood Cancer Program.

## Author contributions

Conceptualization (BG, TG, MPB, LME), Writing – Original Draft (BG), Writing – Review and Editing (BG, TG, EN, SK RJO, SIP, JB, AP, SL, BLG, MNT, CS, SMP, GAG, KAP, MPB, LE) Data Curation (BG, EN SK, RJO, JB, KAP, CS), Formal Analysis (BG, JB, CS), Methodology (BG, SL, BLG, MNT, SMP), Resources (RJO, SIP, AP, GAG, SS, SMP), Funding Acquisition (TG, SMP, GAG, MPB, LME), Supervision (TG, MPB, LME)

## Declaration of Interests

The authors declare no competing interests.

## STAR★Methods

### Key resources table

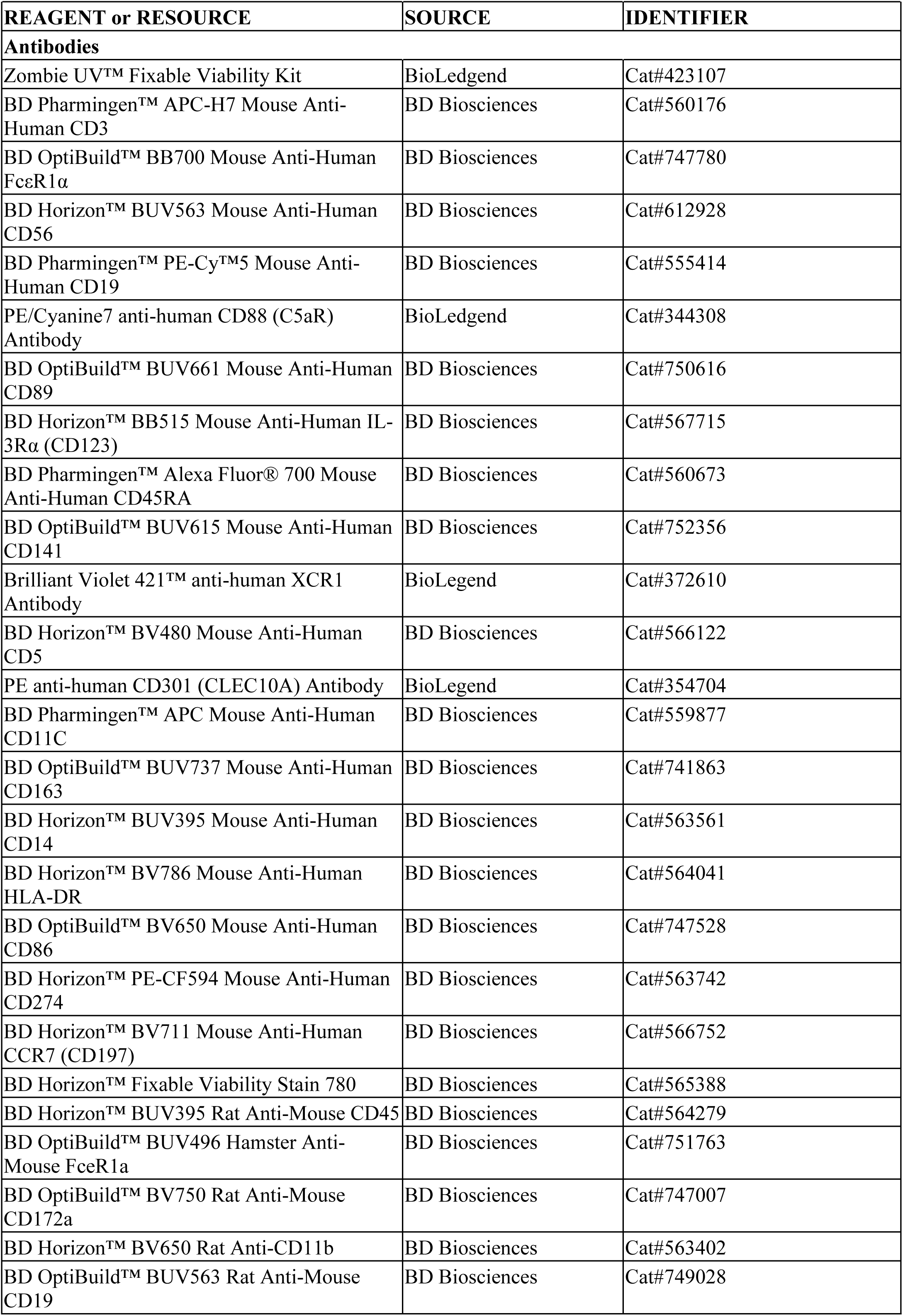
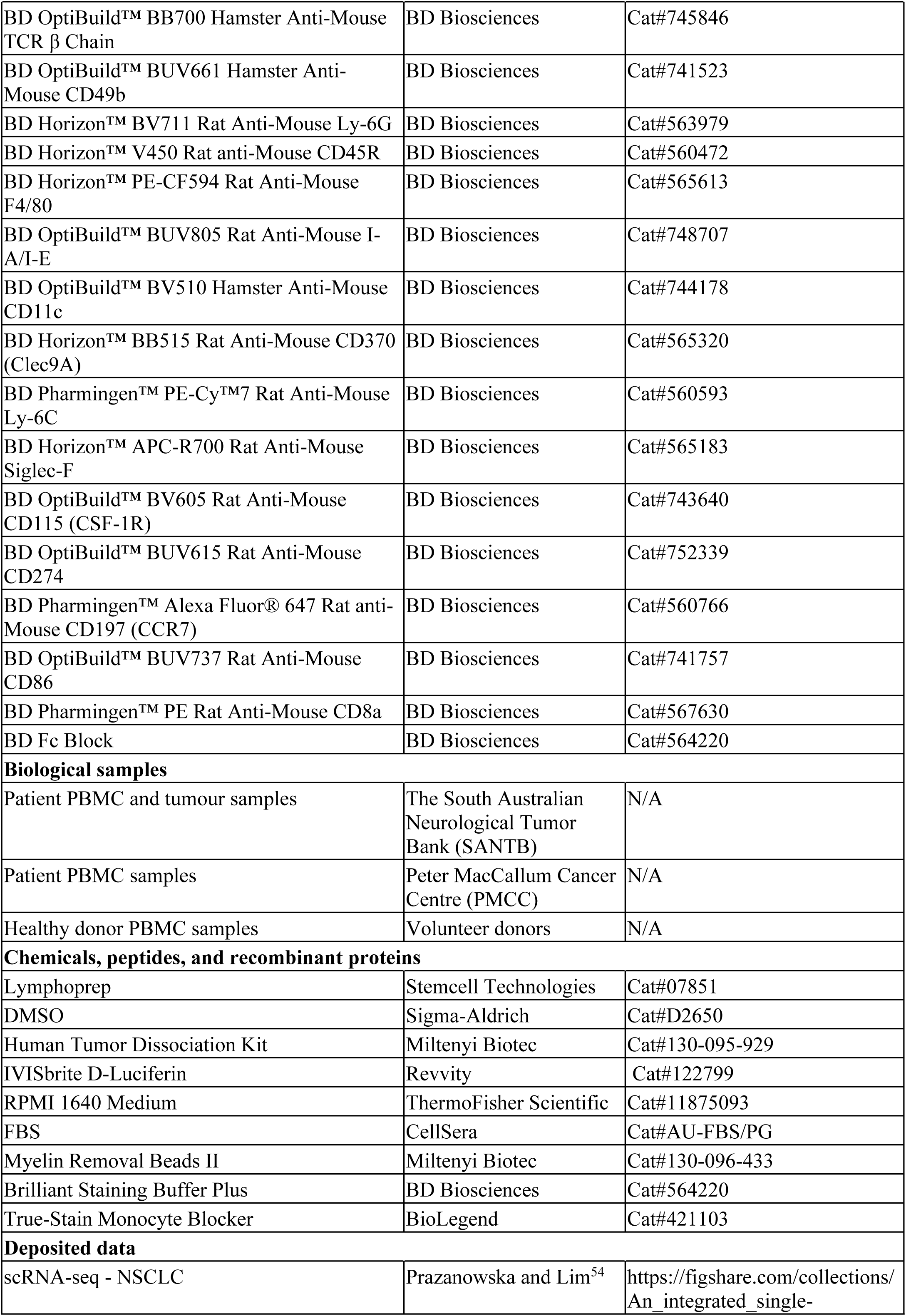
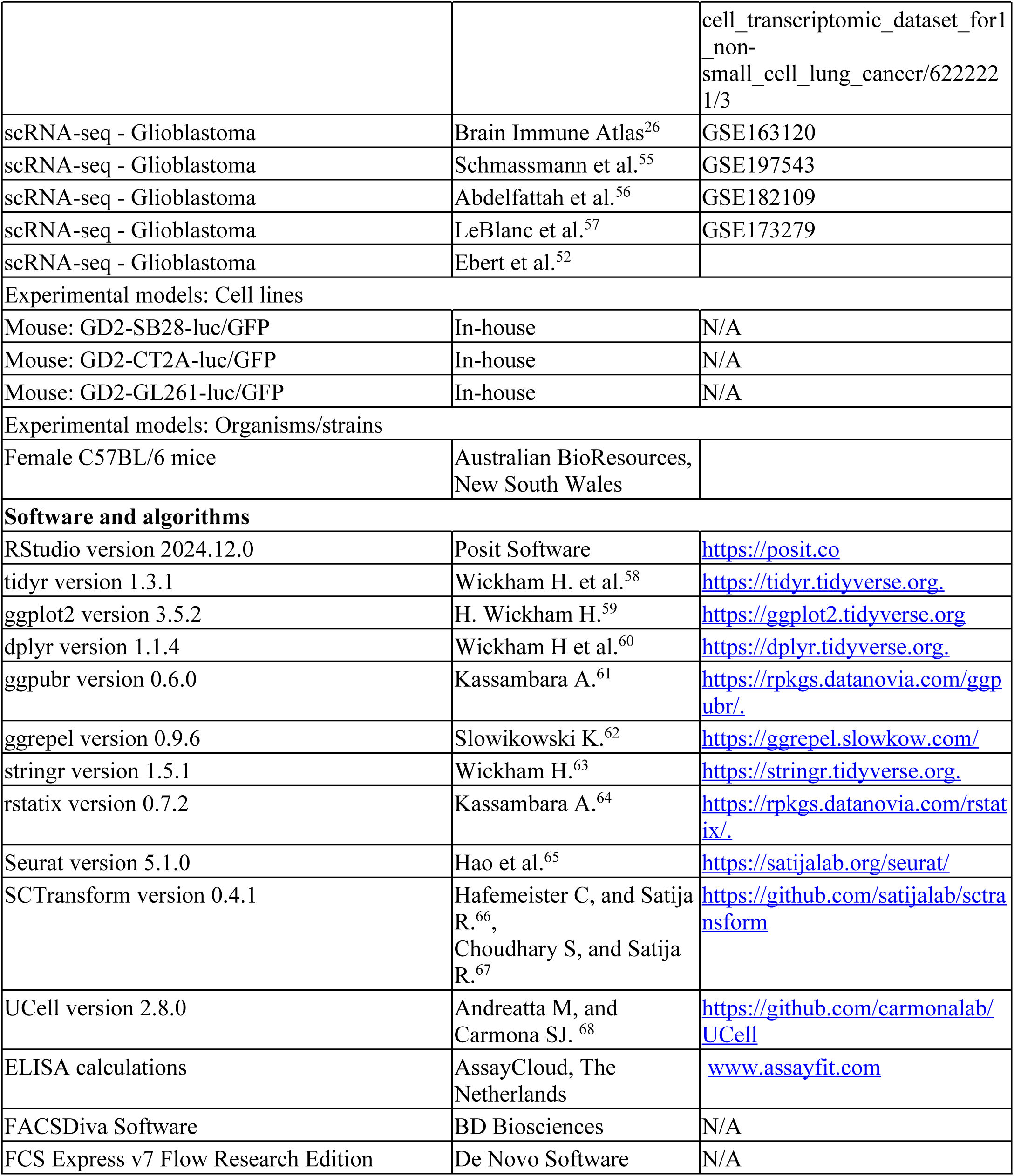

### Experimental model and study participant details

#### Preparation of Whole Blood and Tumor Samples

Blood from volunteer healthy donors was collected into heparin Vacutainers (BD). The South Australian Neurological Tumor Bank (SANTB) and Peter MacCallum Cancer Centre (PMCC) supplied patient blood and tumor samples and associated clinical data. Corticosteroid usage data was collected from medical records. Most brain tumor patients received dexamethasone in line with common practice, although dose and duration of treatment varied considerably (details in Supplementary Table S3). Cancer patients without a brain tumor (non-brain tumor patients) did not routinely receive corticosteroids for management of their cancer, but some had concurrent conditions that required management with corticosteroids. PBMC were purified by density-gradient separation with Lymphoprep (Stemcell Technologies, Cat# 07851). Following density-gradient separation, plasma was collected and stored at −80°C. PBMCs were cryopreserved in 90% fetal bovine serum (FBS) and 10% dimethyl sulfoxide (DMSO) (Sigma-Aldrich, Cat# D2650) for later analysis. Fresh tumor samples were prepared as described previously^52^. Briefly, tumor samples were dissociated to generate a single-cell suspension using the gentleMACS Octo Dissociator in combination with the Human Tumor Dissociation Kit (both Miltenyi Biotec). Following dissociation, cells were filtered through a 70-µm cell strainer and cryopreserved as for PBMC for later analysis.

#### Orthotopic Mouse Brain Tumor Models

A Stoelting motorized stereotactic alignment and injection unit was used to deliver 2 µL of tumor cells over 4 min, 3 mm deep into the right hemisphere of 6–8 weeks old female C57BL/6 mice (Australian BioResources, New South Wales). The following tumor cell lines were used: GD2-SB28-luc/GFP (5×10^3^ cells), GD2-CT2A-luc/GFP (5×10^4^ cells), and GD2-GL261-luc/GFP (1×10^5^ cells). Cell lines were generated from parental lines CT2A (Merck) GL261 (Division of Cancer Treatment and Diagnosis Tumour Repository, NCI) SB28-luc/GFP (DSMZ; Leibniz-InstitutDSMZ Deutsche Sammlung von Mikroorganismen und Zellkulturen GmbH). The cell lines were engineered to express the GD2 antigen by transduction with ecotropic retrovirus encoding murine GD2 and GD2 synthases (MP9956:SFG.GD3synthase-2A-GD2synthase was a gift from Martin Pule Addgene plasmid # 75013; http://n2t.net/addgene:75013^69^; RRID:Addgene_75013; retrovirus generated from the Phoenix-Eco (ATCC Phoenix-ECO CRL-3214) packaging line), with two-rounds of FACS sorting to achieve a >99% GD2+ line. In addition, GD2-CT2A and GD2-GL261 were rendered GFP/F-luciferase positive via lentiviral transduction using pBliv MSCV-copGFP T2A fLuc and FACS sorting on GFP+GD2+ cells. GD2-GL261-luc/GFP were cultured in T-75 flasks in RPMI medium (Thermo Fisher) supplemented with 10% Foetal Bovine Serum (FBS), 1% Glutamax (Thermo Fisher), 1% Non-Essential Amino Acids (NEAA) (Thermo Fisher), and 1% Penicillin-Streptomycin (PenStrep) (Thermo Fisher). GD2-CT2A-luc/GFP were cultured in T-75 flasks in DMEM medium supplemented with 10% FBS, 1% Glutamax, 1% NEAA, and 1% PenStrep. Both cell lines were passaged when they reached 60–80% confluence by detaching cells using TrypLE (Thermo Fisher) and reseded in fresh flasks at 1:3 to 1:6 ratio.

Control mice were sex and age matched, and did not receive surgery or anesthetic. Mice received daily clinical checks and weekly bioluminescence imaging (BLI) to monitor tumor growth. BLI was performed using an IVIS Lumina S5 after intraperitoneal injection of 100 µL of 30 mg/mL IVISbrite D-Luciferin (Revvity, Cat# 122799) in saline. Defined humane endpoints for euthanasia included loss of >15% body weight from starting weight, a body condition score of <2 (under conditioned)^70^, and neurological signs including head-tilt, loss of balance and circling movement. Mice were humanely euthanized with CO_2,_ utilizing the Quietek CO_2_ induction system (Next Advance, USA). When mice reached humane endpoint, spleens were harvested and mechanically dissociated utilizing 40 µm strainers in Roswell Park Memorial Institute (RPMI) 1640 Medium (ThermoFisher Scientific, Cat# 11875093) + 1% FBS (CellSera, Cat# AU-FBS/PG). Samples were centrifuged at 400g for 5 min and resuspended in FBS + 10% DMSO and cryopreserved for later analysis.

#### Flow Cytometry Staining and Analysis

For staining human PBMCs and dissociated tumor specimens, cryopreserved samples were rapidly thawed in a 37°C water bath and transferred to tubes containing 9 mL warm RPMI. For dissociated tumor samples, myelin removal was conducted using magnetic separation beads following the provided protocol (Miltenyi Biotec cat# 130-096-433). Following myelin removal and after 10 min at room temperature (RT) for PBMCs, cells were centrifuged and resuspended in FACS buffer (PBS, 0.01% sodium azide, 0.5% BSA), and Fc receptors were blocked with Human BD Fc Block (BD Biosciences, Cat# 564220) for 10 min at RT. Cells were then incubated for 30 min at RT with cocktails containing Brilliant Staining Buffer Plus (BD Biosciences, Cat# 564220), True-Stain Monocyte Blocker (BioLegend, Cat# 421103), and combinations of the antibodies in Table S4 (human) or Table S5 (mouse). After washing twice in PBS, cells were incubated with a fixable viability stain (Zombie UV, BioLegend Cat #423107) for 30 min at RT, washed in FACS buffer, resuspended in FACS buffer, and stored at 4°C until acquisition on the same day. Mouse samples were transferred to TruCount tubes (BD Biosciences, Cat# 340334) to enable absolute cell numbers to be determined.

All samples were acquired on a BD FACSymphony A5 using FACSDiva Software (BD Biosciences). Analysis was performed using FCS Express v7 Flow Research Edition (De Novo Software).

#### Enzyme-Linked Immunosorbent Assay (ELISA)

ELISAs were conducted on plasma collected during whole blood preparation for the growth factors FLT3L, G-CSF and GM-CSF utilizing DuoSet ELISA kits (R&D Systems Cat# DY308, DY214-05, DY215-05). The assays were performed according to the manufacturer’s instructions. The plates were read on a FLUOstar Omega Plate Reader software version 5.11 RS at 450 nm (BMG LABTECH). The standard curve and results were calculated using the ELISA calculation sheet at www.assayfit.com; all results below the level of detection were replaced with a value of half the limit of detection^53^.

#### Identification of Dendritic Cells in Publicly Available scRNAseq Datasets

Single-cell RNA-sequencing (scRNA-seq) data from NSCLC and glioblastoma patients were obtained from publicly available resources. NSCLC data was obtained from Prazanowska and Lim (https://figshare.com/collections/An_integrated_single-cell_transcriptomic_dataset_for1_non-small_cell_lung_cancer/6222221/3)^54^. Glioblastoma data was obtained from the Brain Immune Atlas (GSE163120)^26^, Schmassmann et al. (GSE197543)^55^, Abdelfattah et al (GSE182109)^56^, LeBlanc et al (GSE173279)^57^, and our previously published dataset^52^.

All data were processed using the Seurat R package (version 5.1.0). Seurat objects were constructed from count matrices and accompanying metadata using the parameters min.cells = 3 and min.features = 200. For lung cancer, datasets were split by study into individual Seurat objects, excluding the data “KU_Loom” due to missing patient metadata and excluding samples with fewer than 750 total cells (GSE153935 and GSE11911). No additional filtering was applied. For GBM data from the Brain Immune Atlas, Seurat objects were filtered to retain cells with <10% mitochondrial gene expression (percent.mt < 10) and fewer than 5500 detected features (nFeature_RNA < 5500). In the dataset GSE197543, periphery and tumor samples were separated from peripheral blood mononuclear cell (PBMC) samples into distinct Seurat objects without further filtering.

#### Data Integration and Annotation

Each Seurat object was split by patient/sample for individual normalization using SCTransform (v0.4.1)^66,67^, with the following parameters: method = glmGamPoi, vst.flavor = “v2”, return.only.var.genes = FALSE, variable.features.n = all features, and min_cells = 0^37^.

A set of 2000 highly variable integration features was selected, supplemented with dendritic cell (DC) marker genes. Integration was performed using SCT normalization and reciprocal PCA (RPCA) reduction. PCA was computed on each dataset using the defined integration features, integration anchors were identified, and datasets were integrated using these anchors.

#### Identification of Immune Cells

Clusters were annotated as immune if more than 70% of cells expressed PTPRC (CD45); others were considered non-immune. Immune cell counts per patient were tabulated across all datasets.

#### Dendritic Cell Subsetting and Annotation

PCA was performed on the integrated object using all features, retaining 200 components. A heatmap of PCA loadings for known DC marker genes across the first 100 components was generated. The number of components to use for UMAP dimensionality reduction was then selected based on the component at which clear separation of DC subtypes first appeared.

UMAP clustering was then performed with the selected dimensions and an empirically determined resolution to optimize DC cluster separation. Lastly, DC-enriched clusters were iteratively subsetted, excluding clusters with high expression of C5AR1, a monocyte marker not typically expressed by cDC2 or DC3 cells^71^.

#### Cell Type Scoring and Quantification

DC subtypes were identified using UCell (version 2.8.0)^68^ to score individual cells based on subtype-specific gene signatures. Scores were smoothed using UCell::SmoothKNN, and cell types were assigned based on the highest smoothed signature score. The number of each DC subtype per patient/sample was then tabulated.

#### Statistical Analysis

Data was graphed and analyzed using RStudio Version 2023.06.1. Details of the packages used are at Table S6.The box plots used throughout display the 3^rd^ quartile (upper line), 1^st^ quartile (lower line), median (centre line), and the maximum range of the data excluding outliers (vertical line). Statistical analysis was conducted in R studio utilizing the Kruskal-Wallis test and the Pairwise Wilcoxon test with Benjamini & Hochberg multiple comparison post-tests. Statistical significance is represented on graphs as * ≤0.05, ** ≤0.01, *** ≤0.001, and **** ≤0.0001.

## Notes

### Competing Interest Statement

The authors have declared no competing interest.

