## Supplementary data for "Tumors Located in the Brain Impair the Frequency and Phenotype of Dendritic Cells in Blood and Tumor"

**(A)**

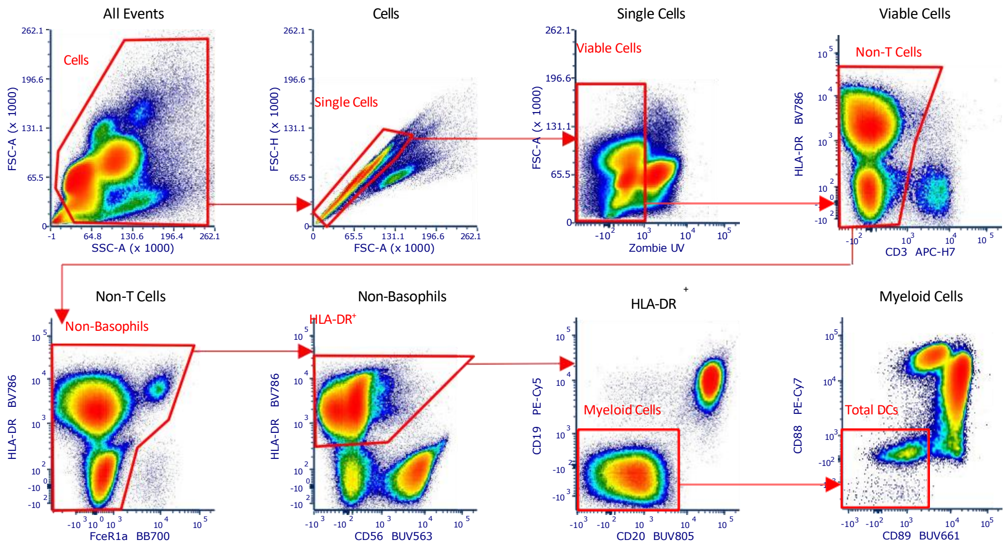


**(B)**

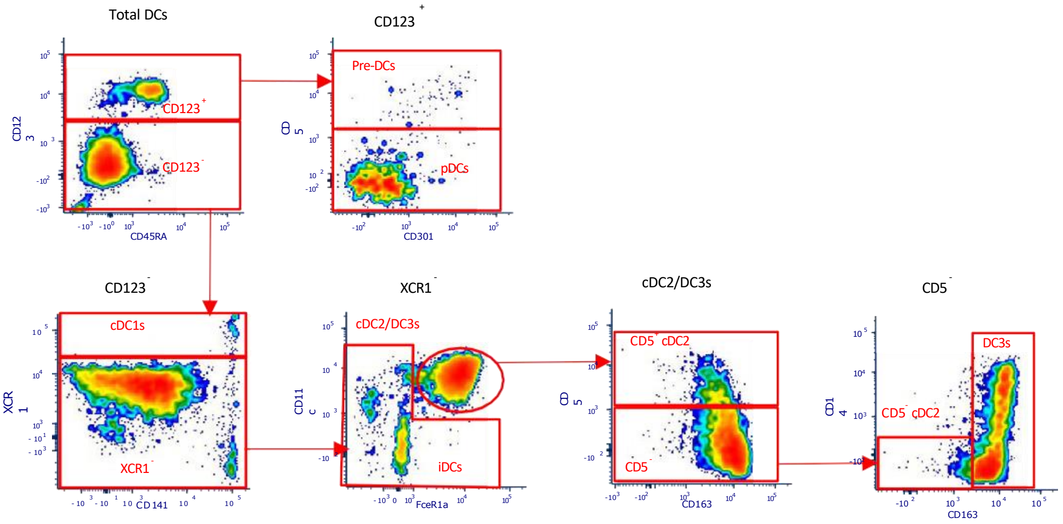


**Figure S1.** (A) Indicative gating strategy of 21 colour flow cytometry panel to determine the total dendritic cells. (B) Indicative gating strategy from total DCs to individual DC subsets adapted from Mair and Leichti (Mair & Liechti 2021).

**Table S1.** Demographics of the healthy donor and patient groups utilized in this study.

| ID | Cohort | Gender (n=) | Median Age (range) | Brain Location | Steroids Before Surgery (time) |
|---|---|---|---|---|---|
| **Summary** | **Healthy Donors** | **Female 47% Male 53% (17)** | **53.94 (29-71)** | | |
| HD1* | Healthy Donors | Female | 44 | N/A | N/A |
| HD1* | Healthy Donors | Female | 47 | N/A | N/A |
| HD2 | Healthy Donors | Female | 39 | N/A | N/A |
| HD5 | Healthy Donors | Female | 29 | N/A | N/A |
| HD9 | Healthy Donors | Female | 63 | N/A | N/A |
| HD10 | Healthy Donors | Female | 56 | N/A | N/A |
| HD12 | Healthy Donors | Female | 65 | N/A | N/A |
| HD14 | Healthy Donors | Female | 70 | N/A | N/A |
| HD3 | Healthy Donors | Male | 45 | N/A | N/A |
| HD4 | Healthy Donors | Male | 35 | N/A | N/A |
| HD6 | Healthy Donors | Male | 39 | N/A | N/A |
| HD7 | Healthy Donors | Male | 48 | N/A | N/A |
| HD8 | Healthy Donors | Male | 60 | N/A | N/A |
| HD11 | Healthy Donors | Male | 71 | N/A | N/A |
| HD13 | Healthy Donors | Male | 71 | N/A | N/A |
| HD15 | Healthy Donors | Male | 67 | N/A | N/A |
| HD16 | Healthy Donors | Male | 68 | N/A | N/A |
| **Summary** | **Primary Glioblastoma** | **Female 43% Male 57% (14)** | **66.14 (46-75)** | | |
| SANTB00501^ | Primary Glioblastoma | Female | 64 | Right Occipital/Temporal | Yes (3 days) |
| SANTB00579 | Primary Glioblastoma | Female | 68 | Left Occipital | Yes (9 days) |
| SANTB00651 | Primary Glioblastoma | Female | 75 | Right Temporal | Yes (2 days) |
| SANTB00679 | Primary Glioblastoma | Female | 72 | Right Parietal | Yes (6 days) |
| SANTB00713 | Primary Glioblastoma | Female | 76 | Left Frontal | Yes (26 days) |
| SANTB00743-1 | Primary Glioblastoma | Female | 53 | Left Frontal | Yes (6 days) |
| SANTB00502 | Primary Glioblastoma | Male | 67 | Right Occipital | Yes (Long term) |
| SANTB00525 | Primary Glioblastoma | Male | 62 | Right Frontal | Yes (5 days) |
| SANTB00598 | Primary Glioblastoma | Male | 75 | Left Frontal | Yes (25 days) |
| SANTB00678 | Primary Glioblastoma | Male | 46 | Right Parietal Inter-axial | Yes (7 days) |
| SANTB00683 | Primary Glioblastoma | Male | 69 | Left Frontal | Yes (7 days) |
| SANTB00690 | Primary Glioblastoma | Male | 67 | Left Temporal | Unk |
| SANTB00706 | Primary Glioblastoma | Male | 64 | Left Temporal | Yes (1 day) |
| SANTB00707 | Primary Glioblastoma | Male | 68 | Right Parietal | Yes (7 days) |
| **Summary** | **Recurrent Glioblastoma** | **Female 63% Male 37% (16)** | **52.25 (33-76)** | | |

| | | | | | |
|---|---|---|---|---|---|
| SANTB00103 | Recurrent Glioblastoma | Female | 55 | Right Temporal | Yes (1 day) |
| SANTB00118-2[#] | Recurrent Glioblastoma | Female | 33 | Left Temporal | Yes (Long term) |
| SANTB00118-3[#] | Recurrent Glioblastoma | Female | 37 | Left Temporal | Yes (3 days) |
| SANTB00501-2^ | Recurrent Glioblastoma | Female | 64 | Right Occipital/Temproal | Yes (15 days) |
| SANTB00485-2[&] | Recurrent Glioblastoma | Female | 51 | Right Frontal | Yes (Long term) |
| SANTB00610 | Recurrent Glioblastoma | Female | 47 | Left Frontal | No |
| SANTB00624 | Recurrent Glioblastoma | Female | 51 | Left Occipital | Yes (1 day) |
| SANTB00544-2 | Recurrent Glioblastoma | Female | 55 | Left Frontal | Yes (1 day) |
| SANTB00485-3[&] | Recurrent Glioblastoma | Female | 53 | Right Frontal | Yes (Long Term) |
| SANTB00486-2 | Recurrent Glioblastoma | Female | 76 | Left Occipital | Yes (1 day) |
| CAREFOR-14 | Recurrent Glioblastoma | Male | 55 | Right Temporal | Unk |
| CAREFOR-15 | Recurrent Glioblastoma | Male | 34 | Right Temporal | Unk |
| SANTB00689 | Recurrent Glioblastoma | Male | 50 | Left Temporal | Yes (2 days) |
| SANTB00663-2 | Recurrent Glioblastoma | Male | 59 | Left Temporal | Yes (Long term) |
| SANTB00496-2 | Recurrent Glioblastoma | Male | 63 | Left Parietal | Yes (1 day) |
| #210 | Recurrent Glioblastoma | Male | 53 | | No |
| **Summary** | **Low Grade Glioma** | **Female 43% Male 57% (7)** | **42.14 (32-61)** | | |
| SANTB00154 | Low Grade Glioma | Female | 33 | Left Frontal | Yes (1 day) |
| SANTB00682 | Low Grade Glioma | Female | 36 | Right Frontal | Yes (6 days) |
| SANTB00702 | Low Grade Glioma | Female | 34 | Right Occipital | Yes (9 days) |
| SANTB00150 | Low Grade Glioma | Male | 43 | Right Frontal | Yes (32 days) |
| SANTB00215 | Low Grade Glioma | Male | 32 | Right Frontal Temporal | Yes (2 days) |
| SANTB00687 | Low Grade Glioma | Male | 56 | Right Frontal | No |
| SANTB00694 | Low Grade Glioma | Male | 61 | Left Frontal | Yes (11 days) |
| **Summary** | **Brain Metastasis** | **Female 23% Male 77% (22)** | **57.32 (30-80)** | | |
| SANTB00555 | Brain Metastasis | Female | 80 | Right Occipital | Yes (1 day) |
| SANTB00260 | Brain Metastasis | Male | 79 | Right Parietal | Yes (9 days) |
| SANTB00503 | Brain Metastasis | Male | 30 | Right Frontal | Yes (Long Term) |
| SANTB00603 | Brain Metastasis | Male | 51 | Right Parietal | Yes (1 day) |
| SANTB00655 | Brain Metastasis | Male | 59 | Left Parietal | Yes (Long Term) |
| SANTB00688 | Brain Metastasis | Male | 73 | Left Frontal | Yes (7 days) |
| SANTB00614 | Brain Metastasis | Male | 69 | Right Parietal | Yes (8 days) |
| SANTB00625 | Brain Metastasis | Male | 81 | Left Frontal | No |

| | | | | | |
|---|---|---|---|---|---|
| SANTB00692 | Brain Metastasis | Male | 61 | Right Frontal | Unk |
| SANTB00693 | Brain Metastasis | Male | 70 | Left Frontal | Yes (3 days) |
| SANTB00725-1 | Brain Metastasis | Male | 79 | Right Frontal | Yes (2 days) |
| #303 | Brain Metastasis | Male | 45 | Right Frontal | No |
| #303(2) | Brain Metastasis | Male | 49 | Right Frontal | No |
| #304 | Brain Metastasis | Male | 40 | Left Parietal and Left Occipital | No |
| PMC0081-1 | Brain Metastasis | Female | 45 | Left Frontal | No |
| BIO-005 | Brain Metastasis | Female | 45 | Left Temporal | No |
| BIO-091 | Brain Metastasis | Female | 39 | Posterior Lobe | No |
| PMC0081-2 | Brain Metastasis | Female | 45 | Left Frontal | No |
| BIO-120 | Brain Metastasis | Male | 53 | Left Cerebellar | No |
| BIO-140 | Brain Metastasis | Male | 56 | Left Frontal | No |
| BIO-222 | Brain Metastasis | Male | 50 | Right Temporal | No |
| CHA020 | Brain Metastasis | Male | 62 | Left occipital | No |
| **Summary** | **Type of Primary Cancer** | **Female 59% Male 41% (17)** | **66.88 (50-84)** | | |
| Non-brain tumor 1 | Melanoma | Female | 75 | None | No |
| Non-brain tumor 2 | Melanoma | Female | 58 | None | No |
| Non-brain tumor 4 | Melanoma | Female | 50 | None | No |
| Non-brain tumor 6 | Melanoma | Female | 77 | None | No |
| Non-brain tumor 7 | Lung Cancer | Female | 66 | None | Yes (Long Term) |
| Non-brain tumor 8 | Lung Cancer | Female | 62 | None | Yes (Long Term) |
| Non-brain tumor 9 | Lung Cancer | Female | 71 | None | No |
| Non-brain tumor 11 | Lung Cancer | Female | 62 | None | No |
| Non-brain tumor 12 | Lung Cancer | Female | 69 | None | No |
| Non-brain tumor 13 | Melanoma | Female | 73 | None | Yes (9 days) |
| Non-brain tumor 3 | Melanoma | Male | 66 | None | No |
| Non-brain tumor 5 | Melanoma | Male | 78 | None | No |
| Non-brain tumor 10 | Lung Cancer | Male | 80 | None | No |
| Non-brain tumor 14 | Melanoma | Male | 81 | None | Yes (4 days) |
| Non-brain tumor 15 | Melanoma | Male | 74 | None | Yes (Long Term) |
| Non-brain tumor 16 | Melanoma | Male | 84 | None | Yes (Long Term) |
| Non-brain tumor 17 | Melanoma | Male | 71 | None | No |

Notes:
* Two individual donations
#, ^,& Are the same patients
Steroids Before Surgery Yes (1 day) is for the day of surgery only

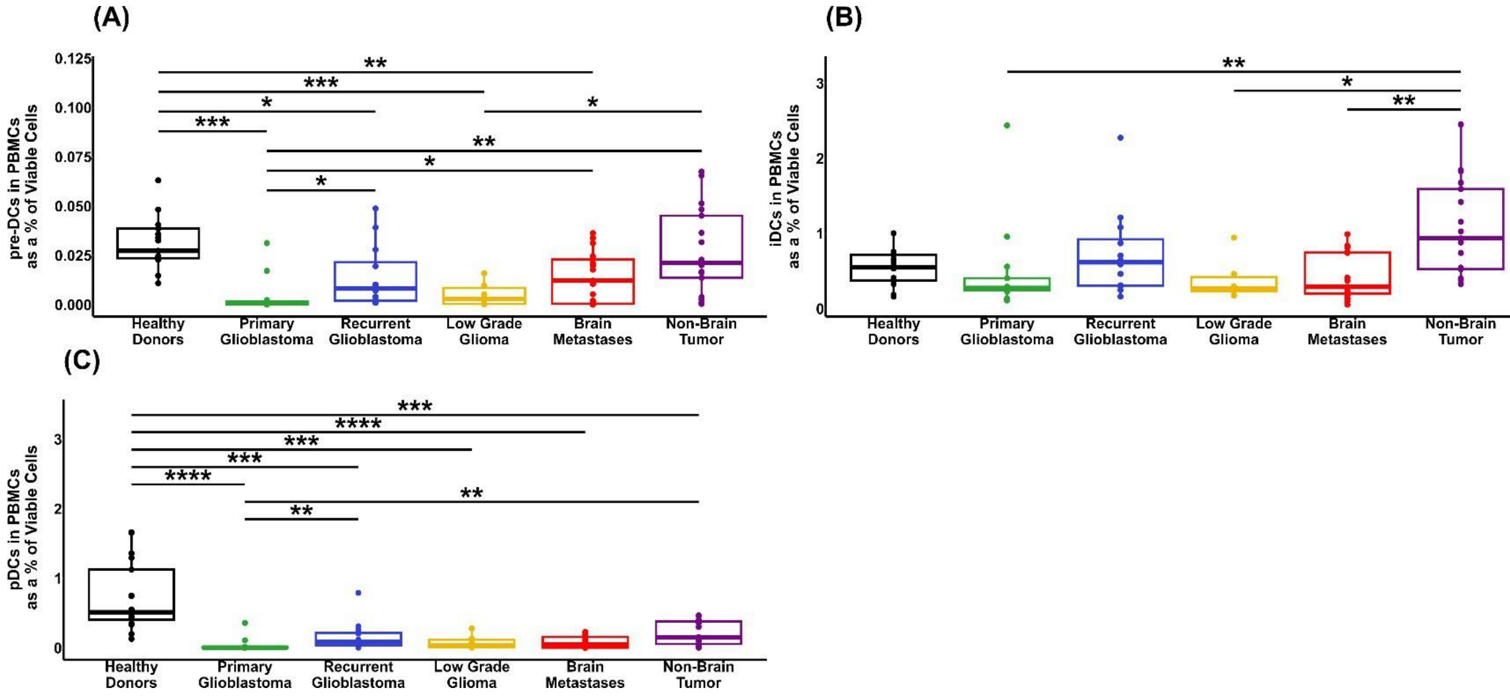


**Figure S2.** (A) Quantification of the percentages of pre-DCs as a percentage of the total viable cells from PBMCs. (B) Quantification of the percentages of iDCs as a percentage of the total viable cells from PBMCs. (C) Quantification of the percentages of pDCs as a percentage of the total viable cells from PBMCs.

P values *p<0.05, **P<0.01, ***p<0.001, ****p<0.0001

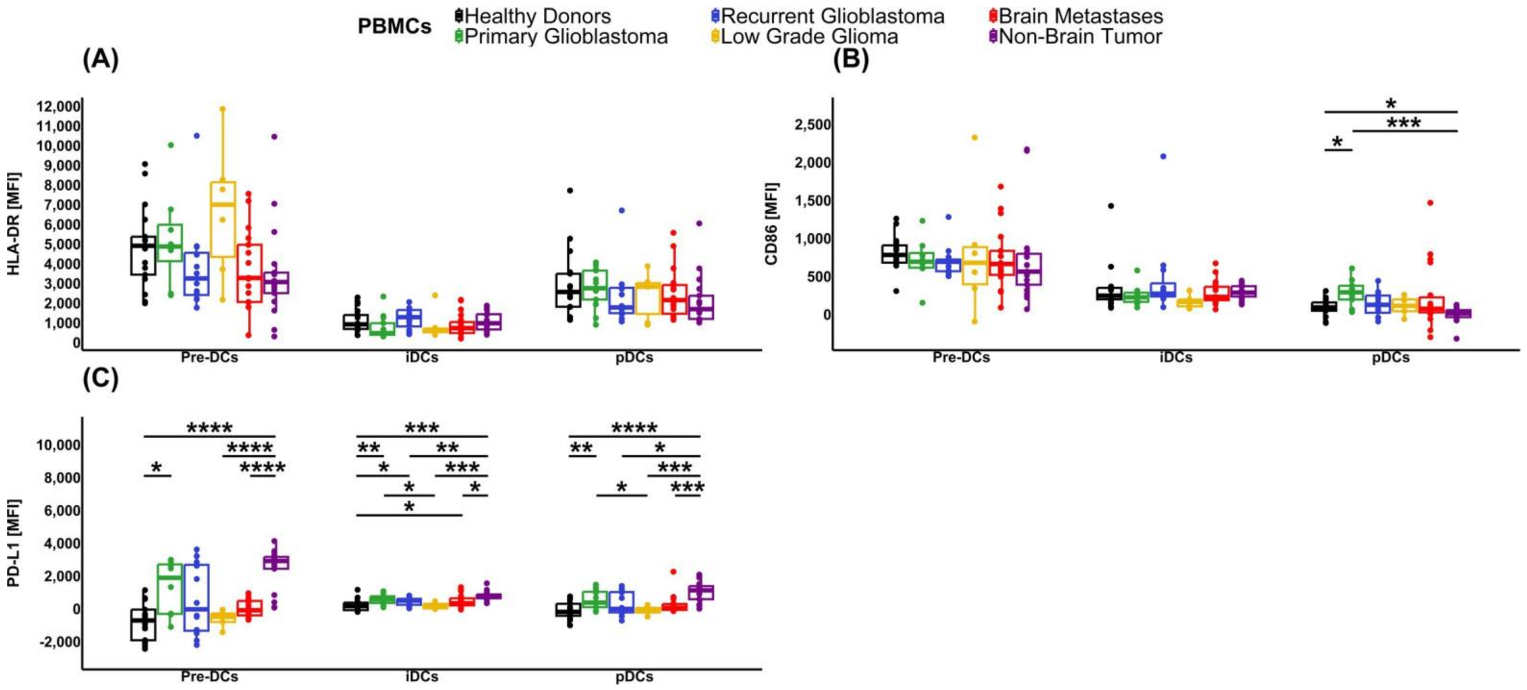


**Figure S3** Quantification of pre-DCs, iDCs, and pDCs from PBMCs for (A) the MFI of HLA-DR. (B) the MFI of CD86, and (C) the MFI of PD-L1.

P values *p<0.05, **P<0.01, ***p<0.001, ****p<0.0001

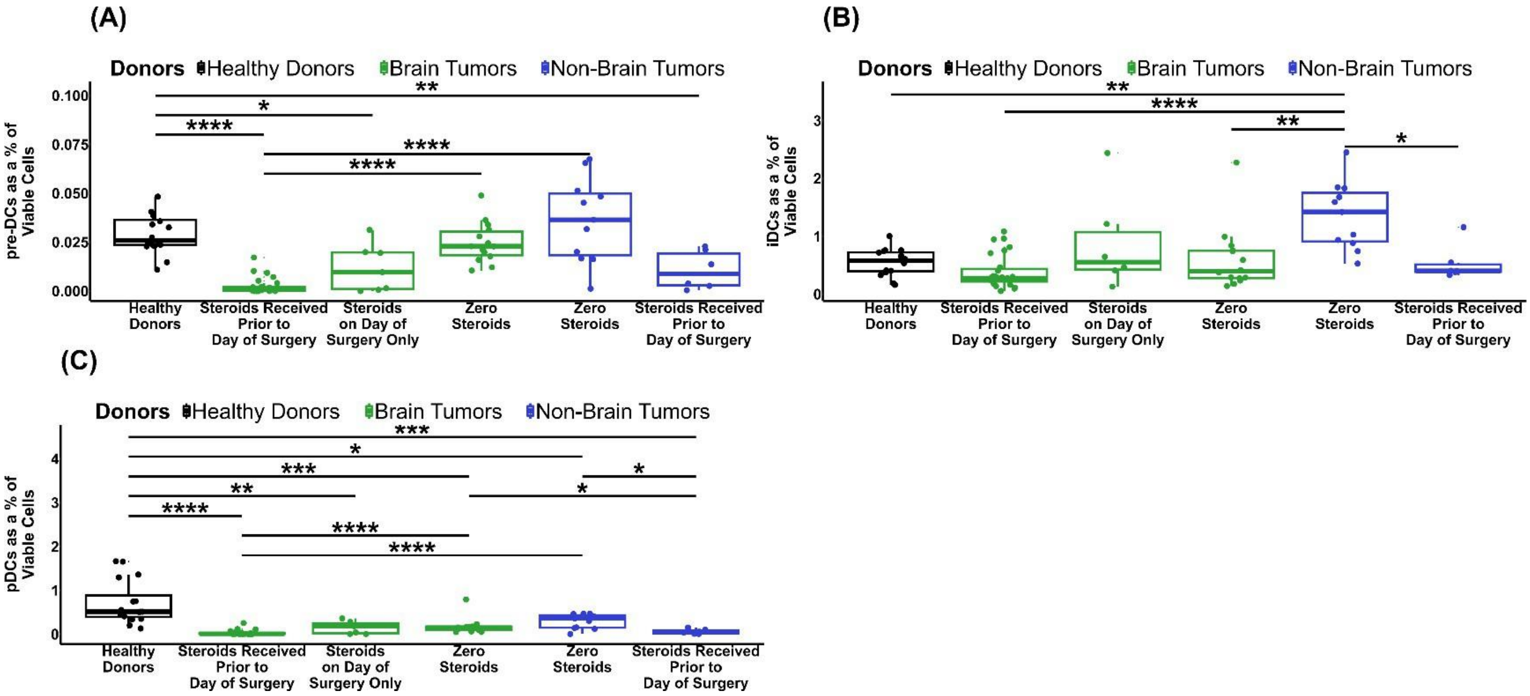


**Figure S4.** Steroid usage in combined brain tumour patients and non-brain tumour patients in (A) pre-DCs, (B) iDCs, and (C) pDCs.

P values *p<0.05, **P<0.01, ***p<0.001, ****p<0.0001

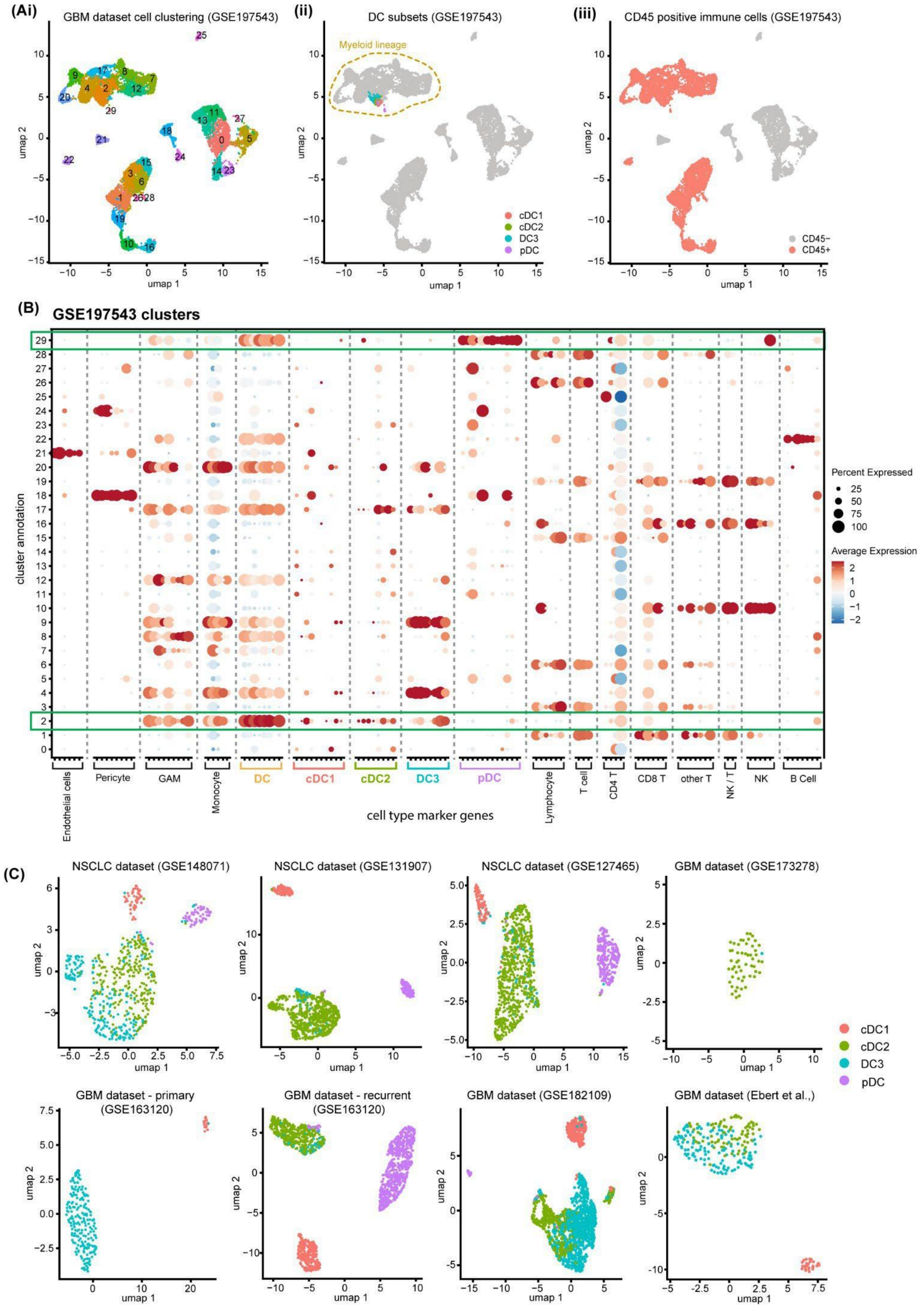


**Figure S5** (A) UMAP plots illustrating (i) total cell population clustering, (ii) localisation of dendritic cells (DCs) within the myeloid lineage cluster, and (iii) annotation of $CD45^{+}$ immune cells in a GBM scRNA-seq dataset (GSE197543). (B) Expression of cell type markers across cell clusters in the GBM dataset (GSE197543); green boxes indicate enrichment of DC subset marker genes in clusters 2 and 29. (C) UMAP plots showing DC subsets in all NSCLC and GBM datasets (datasets GSE197543 and GSE136246 shown in Figure 5).

**Table S2.** List of broad cell type and DC subset-specific marker genes utilized in the scRNA-seq analysis

| Cell Type | Markers | | | |
|---|---|---|---|---|
| Endothelial cells | VWF | CLDN5 | CD34 | |
| | APOLD1 | TIE1 | CDH5 | |
| Pericyte | PDGFRB | COL4A2 | COL4A1 | ACTA2 |
| | MCAM | CSPG4 | FN1 | THY1 |
| Glioma Accosiated Macrphage | AIF1 | CSF1R | ITGAX | TREM2 |
| | FCER1G | GPR34 | LAPTM5 | CD83 |
| | RGS10 | | | |
| Lymphocyte | CD2 | CD6 | IL7R | CD96 |
| | CD247 | LTB | | |
| Monocyte/Macrophage | C5AR1 | | | |
| Dendritic Cell | CD74 | HLA-DRB1 | HLA-DQA1 | CST3 |
| | HLA-DPA1 | HLA-DPB1 | HLA-DRA | HLA-DQB1 |
| cDC1 | CADM1 | CLNK | C1orf54 | RAB32 |
| | XCR1 | WFDC21P | IDO1 | BATF3 |
| | CLEC9A | DNASE1L3 | LGALS2 | FLT3 |
| cDC2 | CD1C | CD1E | CD86 | RALA |
| | FCER1A | CLEC4A | C15orf48 | PKIB |
| | CLEC10A | RALA | | |
| DC3 | VCAN | S100A9 | CD36 | LYZ |
| | S100A8 | FCN1 | CD163 | CD14 |
| pDC | IL3RA | ASIP | TPM2 | SPIB |
| | BCL11A | PTCRA | MZB1 | ITM2C |
| | TCF4 | LRRC26 | SMPD3 | SERPINF1 |
| | LILRA4 | | | |
| Monocyte | CTSS | NEAT1 | PSAP | SERPINA1 |
| T Cells | CD3D | CD3G | TRAC | |
| CD4 + T Cells | MAL | CD4 | LDHB | TPT1 |
| CD8+ Tcells | CD8B | CD8A | HCST | CD3E |
| | LINC02446 | CTSW | | |
| Other T Cells | TRDC | GZMK | KLRB1 | GZMA |
| | TRGC2 | LYAR | KLRG1 | |
| NK T Cells | NKG7 | CST7 | | |
| NK Cells | KLRD1 | GNLY | PRF1 | GZMB |
| | KLRF1 | | | |
| B Cells | CD791 | RALGPS2 | CD79B | IGHM |
| | MS4A1 | BANK1 | TNFRSF13C | MEF2C |

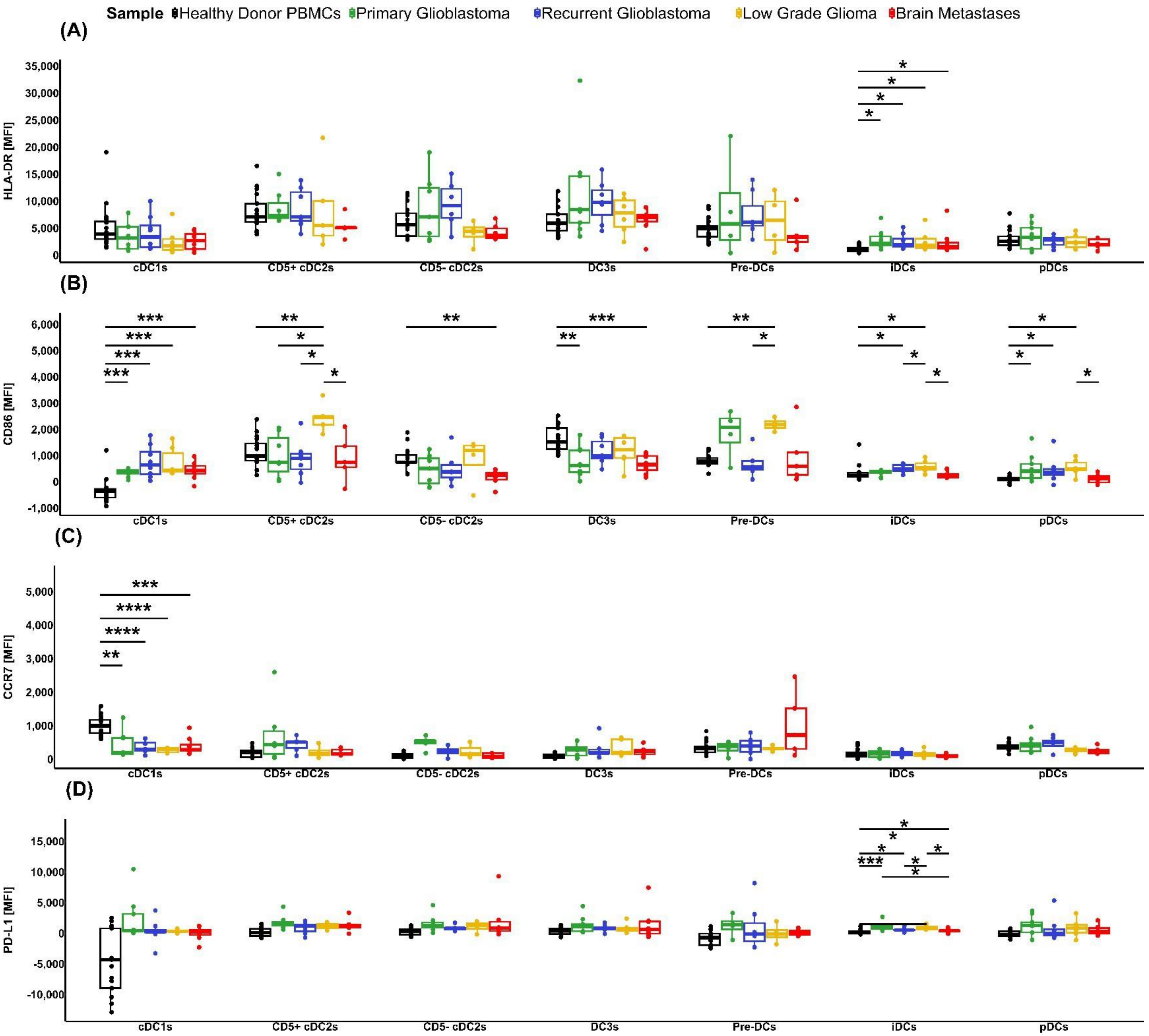


**Figure S6.** Quantification of DC subsets in patient tumor samples compared to healthy donor PBMCs for (A) the MFI of HLA-DR. (B) the MFI of CD86, (C) MFI of CCR7 and (D) the MFI of PD-L1. P values *p<0.05, **P<0.01, ***p<0.001, ****p<0.0001

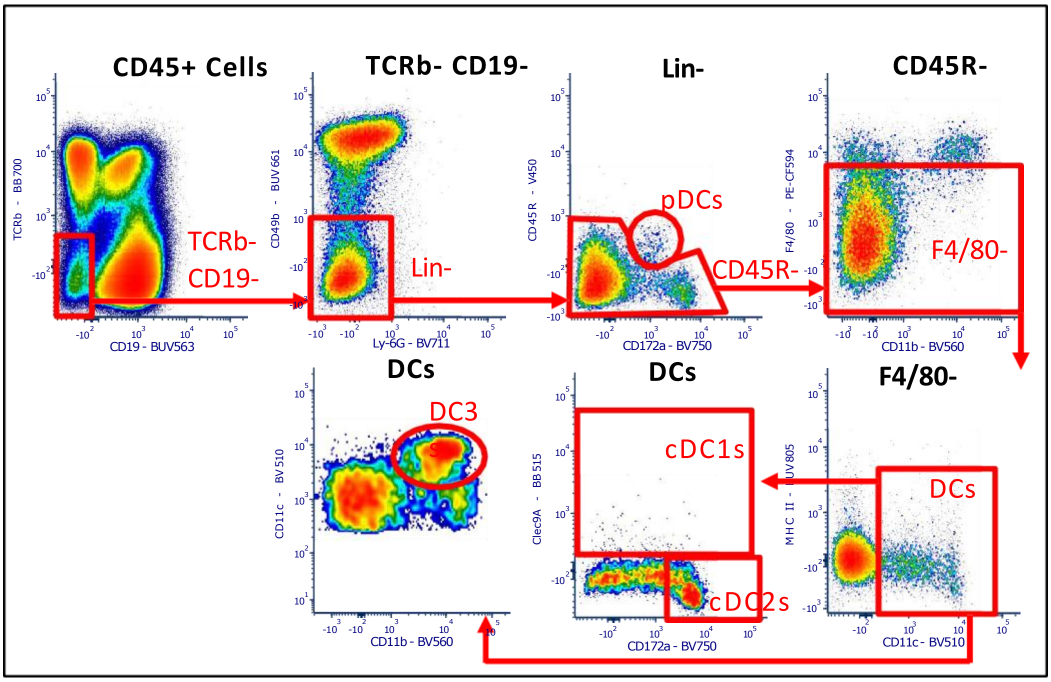


**Figure S7.** Indicative gating strategy for C57BL/6 mouse to identify the murine dendritic cells and subsets

**Table S3** . Corticosteroid usage data for patients with brain tumours

| ID | Gender | Age | Pathological diagnosis | Date of surgery | Steroid date commenced | Steroid | Steroid dosage |
|---|---|---|---|---|---|---|---|
| SANTB00118-2 | Female | 33 | WHO Grade IV | 16/10/2017 | Pre-admission | 1mg Dexamethasone | 4mg x 1 times day for surgery |
| SANTB00118-3 | Female | 37 | WHO Grade IV | 5/05/2021 | 3/05/2021 | Nil pre-admission | 8mg x 2 times for surgery |
| SANTB00150 | Male | 43 | Grade III anaplastic astrocytoma | 11/10/2017 | 28/09/2017 | 2mg Dexamethasone | 4mg x 3 times a day on 6/10/17 |
| SANTB00154 | Female | 33 | Grade III anaplastic oligodendroglioma | 25/10/2017 | 25/10/2017 | Nil pre-admission | 4mg x 4 times a day for surgery |
| SANTB00215 | Male | 32 | High-grade astrocytoma | 25/07/2018 | 23/07/2018 | Nil pre-admission | 4mg x 4 times a day for surgery |
| SANTB00260 | Male | 79 | Metastatic squamous cell carcinoma | 30/01/2019 | 22/01/2019 | Nil pre-admission | 4mg x 4 times a day for surgery |
| SANTB00485-2 | Female | 51 | WHO Grade IV | 8/11/2021 | 1/11/2020 | 0.5mg x 1 long-term Dexamethasone | 4mg x 3 for surgery |
| SANTB00485-3 | Female | 53 | WHO Grade IV | 22/05/2023 | pre admission | 0.5mg x 1 long-term Dexamethasone | 0.5mg x 1 times a day for surgery |
| SANTB00496-2 | Male | 63 | WHO Grade IV | 5/04/2023 | 5/04/2023 | Nil pre-admission | 4mg x 3 times a day for surgery |
| SANTB00501 | Female | 64 | WHO Grade IV | 16/12/2020 | 14/12/2020 | Nil pre-admission | 8mg x 2 times on day of surgery |
| SANTB00501-2 | Female | 64 | WHO Grade IV | 8/05/2021 | 24/03/2021 | unsure | unsure |
| SANTB00502 | Male | 67 | WHO Grade IV | 4/01/2021 | pre admission | 2mg Dexamethasone continuing | 8mg x 2 times on day surgery |
| SANTB00503 | Male | 30 | Metastatic sarcoma | 4/01/2021 | pre admission | 2mg Dexamethasone | 8mg x 2 times on day surgery |
| SANTB00525 | Male | 62 | WHO Grade IV | 25/03/2021 | 21/03/2021 | Nil pre-admission | 8mg x 2 times on day surgery |
| SANTB00546 | Female | 41 | WHO Grade IV | 28/04/2021 | 24/04/2021 | Nil pre-admission | 8mg x 2 times for surgery |
| SANTB00555 | Female | 80 | Metastatic carcinoma | 18/05/2021 | 18/05/2021 | Nil pre-admission | 4mg x 4 times day on 18/5/21 |
| SANTB00579 | Female | 68 | WHO Grade IV | 13/08/2021 | 5/08/2021 | Dexamethasone | 4mg x 4 times a day for surgery |
| SANTB00598 | Male | 75 | WHO Grade IV | 22/10/2021 | 28/09/2021 | Dexamethasone | 4mg x 4 times a day for surgery |
| SANTB00603 | Male | 51 | High-grade B-cell lymphoma | 3/11/2021 | surgery only | Nil pre-admission | 4mg x 4 times a day for surgery |
| SANTB00610 | Female | 47 | WHO Grade IV | 24/11/2021 | nil | nil | nil |
| SANTB00624 | Female | 51 | WHO Grade IV | 21/03/2022 | surgery only | nil | 4mg x 4 times a day for surgery |
| SANTB00651 | Female | 75 | WHO Grade IV | 18/07/2022 | 16/06/2022 | nil | 4mg x 4 times a day for surgery |
| SANTB00663-2 | Male | 59 | WHO Grade IV | 21/03/2023 | pre admission | 0.25mg x1 long-term Dexamethasone | 4mg x 4 times day for surgery |
| SANTB00679 | Female | 72 | WHO Grade IV | 9/01/2023 | 4/01/2023 | nil pre-admission | 4mg x 3 times a day for surgery |
| SANTB00682 | Female | 36 | Oligodendroglioma WHO Grade II | 24/01/2023 | 19/01/2023 | Nil pre-admission | 8mg x 2 times per day for surgery |

| | | | | | | | |
|---|---|---|---|---|---|---|---|
| SANTB00683 | Male | 69 | WHO Grade IV | 31/01/2023 | 24/01/2023 | nil pre-admission | 4mg x 3 times a day for surgery |
| SANTB00687 | Male | 56 | Oligodendroglioma WHO Grade II | 28/02/2023 | nil | nil pre-admission | nil |
| SANTB00688 | Male | 73 | Metastatic melanoma | 7/03/2023 | 1/03/2023 | nil pre-admission | 4mg x 4 times day for surgery |
| SANTB00689 | Male | 50 | WHO Grade IV | 8/03/2023 | 7/03/2023 | nil pre-admission | 8mg x 2 times per day for surgery |
| SANTB00692 | Male | 61 | Metastatic clear cell renal cell carcinoma | 16/03/2023 | unsure | nil pre-admission | 4mg x 3 times a day for surgery |
| SANTB00693 | Male | 70 | Metastatic poorly differentiated carcinoma | 20/03/2023 | 23/03/2023 | nil pre-admission | 4mg x 3 times a day for surgery |
| SANTB00694 | Male | 61 | Anaplastic oligodendroglioma | 29/03/2023 | 19/03/2023 | Nil pre-admission | 4mg x 2 times a day for surgery |
| SANTB00706 | Male | 64 | WHO Grade IV | 9/05/2023 | 9/05/2023 | Nil pre-admission | 4mg x 4 times day for surgery |
| SANTB00614 | Male | 69 | Metastatic melanoma | 12/01/2022 | 5/01/2022 | Nil pre-admission | 8mg BD |
| SANTB00707 | Male | 68 | WHO Grade IV | 19/05/2023 | 13/05/2023 | 4mg x 4 per day Dexamethasone | 4mg x 4 per day for surgery |
| SANTB00713 | Female | 76 | WHO Grade IV | 5/06/2023 | 10/5/23 (4 days) 4mg 2x day to wean | nil on admission | 4mg x 4 per day for surgery |
| SANTB00486-2 | Female | 76 | WHO Grade IV | 31/07/2023 | 31/07/2023 | nil on admission | 4mg x 2 per day for surgery |
| SANTB00743-1 | Female | 53 | WHO Grade IV | 12/09/2023 | 6/09/2023 | nil pre-admission | 4mg x 4 per day for surgery |
| NonCNS 1 | Female | 75 | Metastatic melanoma | 25/05/2021 | | nil pre-admission | |
| NonCNS 2 | Female | 53 | Metastatic melanoma | 30/07/2021 | | nil pre-admission | |
| NonCNS 3 | Male | 66 | Metastatic melanoma cervical LN met | 19/08/2022 | | nil pre-admission | |
| NonCNS 4 | Female | 50 | Metastatic melanoma | Jul-21 | | prednisolone | 10mg daily |
| NonCNS 5 | Male | 78 | Metastatic melanoma | Feb-21 | | nil pre-admission | |
| NonCNS 6 | Male | 77 | Metastatic melanoma | Not specified | | nil pre-admission | |
| NonCNS 7 | Female | 66 | Recurrent NSCLC | Not specified | | Prednisolone | 37.5mg-10mg daily |
| NonCNS 8 | Female | 62 | Metastatic NSCLC | Not specified | 18/12/2023 | beclometasone - 100microgram-6microgram-10microgram | Twice daily |
| NonCNS 9 | Female | 71 | Metastatic NSCLC | Not specified | | dexamethasone | dexamethasone 2mg daily |
| NonCNS 10 | Male | 80 | NSCLC | Not specified | | nil pre-admission | |
| NonCNS 11 | Female | 62 | Metastatic NSCLC | Not specified | | nil pre-admission | |
| NonCNS 12 | Female | 69 | Metastatic NSCLC | 15/03/2023 | | prednisolone | 5mg daily |
| NonCNS 13 | Female | 73 | Metastatic melanoma | Not specified | | dexamethasone | 4mg twice daily |
| NonCNS 14 | Male | 81 | Metastatic melanoma | Not specified | | prednisolone | 5mg daily |

| | | | | | | | |
|---|---|---|---|---|---|---|---|
| NonCNS 15 | Male | 74 | Metastatic melanoma | Not specified | | methylprednisolone | 2mg/kg |
| NonCNS 16 | Male | 84 | Metastatic melanoma | Not specified | | prednisolone | not specified |
| NonCNS 17 | Male | 71 | Metastatic melanoma | Not specified | | dexamethasone | 8mg daily |

**Table S4.** Antibodies that were utilized for staining human PBMCs and tumour samples for the identification of DC and subsets.

| Antibody | Conjugate | Clone | Supplier | Catalogue # |
|---|---|---|---|---|
| Viability | Zombie UV | | BioLedgend | 423107 |
| CD3 | APC-H7 | SK7 | BD Biosciences | 560176 |
| FcεR1α | BB700 | AER-37 | BD Biosciences | 747780 |
| CD56 | BUV563 | NCAM16.2 | BD Biosciences | 612928 |
| CD19 | PE-Cy5 | HIB19 | BD Biosciences | 555414 |
| CD88 | PE-Cy7 | S5/1 | BioLedgend | 344308 |
| CD89 | BUV661 | A59 | BD Biosciences | 750616 |
| IL-3Rα (CD123) | BB515 | 6H6 | BD Biosciences | 567715 |
| CD45RA | AF700 | HI100 | BD Biosciences | 560673 |
| CD141 | BUV615 | 1A4 | BD Biosciences | 752356 |
| XCR1 | BV421 | S15064E | BioLedgend | 372610 |
| CD5 | BV480 | UCHT2 | BD Biosciences | 566122 |
| CD301 (CLEC10A) | PE | H037G3 | BioLedend | 354704 |
| CD11c | APC | B-ly6 | BD Biosciences | 559877 |
| CD163 | BUV737 | GHI/61 | BD Biosciences | 741863 |
| CD14 | BUV395 | MφP9 | BD Biosciences | 563561 |
| HLA-DR | BV786 | G46-6 | BD Biosciences | 564041 |
| CD86 | BV650 | BU63 | BD Biosciences | 747528 |
| CD274 (PD-L1) | PE-CF594 | MIH1 | BD Biosciences | 563742 |
| CCR7 (CD197) | BV711 | 2-L1-A | BD Biosciences | 566752 |

**Table S5.** Antibodies that were utilized for staining murine tissue samples for the identification of DC and subsets.

| Antibody | Conjugate | Clone | Supplier | Catalogue # |
|---|---|---|---|---|
| Viability | FVS 780 | | BD Biosciences | 565388 |
| CD45 | BUV395 | 30-F11 | BD Biosciences | 564279 |
| FcεR1α | BUV496 | MAR-1 | BD Biosciences | 751763 |
| CD172a | BV750 | P84 | BD Biosciences | 747007 |
| CD11b | BV650 | M1/70 | BD Biosciences | 563402 |
| CD19 | BUV563 | 1D3 | BD Biosciences | 749028 |
| TCR b | BB700 | H57-597 | BD Biosciences | 745846 |
| CD49b | BUV661 | HMα2 | BD Biosciences | 741523 |
| Ly-6G | BV711 | 1A8 | BD Biosciences | 563979 |
| CD45R | V450 | MAR-1 | BD Biosciences | 751763 |
| F4/80 | PE-CF594 | T45-2342 | BD Biosciences | 565613 |
| I-A/I-E (MHC II) | BUV805 | 2G9 | BD Biosciences | 748707 |
| CD11c | BV510 | N418 | BD Biosciences | 744178 |
| Clec9A (CD370) | BB515 | 10B4 | BD Biosciences | 565320 |
| Ly-6C | PE-Cy7 | AL-21 | BD Biosciences | 560593 |
| Siglec-F | APC-R700 | E50-2440 | BD Biosciences | 565183 |
| CD115 | BV605 | T38-320 | BD Biosciences | 743640 |
| PD-L1 (CD274) | BUV615 | MIH5 | BD Biosciences | 752339 |
| CCR7 (CD197) | AF647 | 4B12 | BD Biosciences | 560766 |
| CD86 | BUV737 | PO3 | BD Biosciences | 741757 |
| CD8a | PE | 5H10-1 | BD Biosciences | 567630 |

**Table S6**. Packages utilized in RStudio**.**

| Package | Version | Reference |
|---|---|---|
| tidyr | 1.3.1 | Wickham H, Vaughan D, Girlich M (2024). *tidyr: Tidy Messy Data*. R package version 1.3.1, https://github.com/tidyverse/tidyr, https://tidyr.tidyverse.org. |
| ggplot2 | 3.5.2 | H. Wickham. ggplot2: Elegant Graphics for Data Analysis. Springer-Verlag New York, 2016. |
| dplyr | 1.1.4 | Wickham H, François R, Henry L, Müller K, Vaughan D (2023). *dplyr: A Grammar of Data Manipulation*. R package version 1.1.4, https://github.com/tidyverse/dplyr, https://dplyr.tidyverse.org. |
| ggpubr | 0.6.0 | Kassambara A (2023). *ggpubr: 'ggplot2' Based Publication Ready Plots*. R package version 0.6.0, https://rpkgs.datanovia.com/ggpubr/. |
| ggrepel | 0.9.6 | Slowikowski K (2024). *ggrepel: Automatically Position Non-Overlapping Text Labels with 'ggplot2'*. https://ggrepel.slowkow.com/, https://github.com/slowkow/ggrepel. |
| stringr | 1.5.1 | Wickham H (2023). *stringr: Simple, Consistent Wrappers for Common String Operations*. R package version 1.5.1, https://github.com/tidyverse/stringr, https://stringr.tidyverse.org. |
| rstatix | 0.7.2 | Kassambara A (2023). *rstatix: Pipe-Friendly Framework for Basic Statistical Tests*. R package version 0.7.2, https://rpkgs.datanovia.com/rstatix/. |
